# Stress granules are shock absorbers that prevent excessive innate immune responses to dsRNA

**DOI:** 10.1101/2021.04.26.441141

**Authors:** Max Paget, Cristhian Cadena, Sadeem Ahmad, Hai-Tao Wang, Tristan X. Jordan, Ehyun Kim, Beechui Koo, Shawn M. Lyons, Pavel Ivanov, Benjamin tenOever, Xin Mu, Sun Hur

## Abstract

Proper defense against microbial infection depends on the controlled activation of the immune system. This is particularly important for the RIG-I-like receptors (RLRs), which recognize viral dsRNA and initiate antiviral innate immune responses with the potential of triggering systemic inflammation and immunopathology. Here we show that stress granules (SGs), molecular condensates that form in response to various stresses including viral dsRNA, play key roles in controlled activation of RLR signaling. Without the SG nucleators G3BP1/2 and UBAP2L, dsRNA triggers excessive inflammation and immune-mediated apoptosis. In addition to exogenous dsRNA, we find that host-derived dsRNA generated in response to ADAR1 deficiency is also controlled by SG biology. Intriguingly, SGs can function beyond immune control by suppressing viral replication independent of the RLR pathway. These observations thus highlight the multi-functional nature of SGs as cellular “shock absorbers” that converge on protecting cell homeostasis–by dampening both toxic immune response and viral replication.

## Introduction

Detection of foreign nucleic acids is central to innate immune defense in all kingdoms of life (tenOever, 2016). Double-stranded RNA (dsRNA) is one such foreign nucleic acid that triggers a wide range of innate immune responses in vertebrates. It has long been thought that dsRNAs are produced only during viral infection as a result of RNA-dependent RNA polymerization of the viral RNA genome or convergent bi-directional transcription of the viral DNA genome (Son et al., 2015; Weber et al., 2006). However, recent studies suggest that dsRNA can also be produced from many dysregulated cellular processes, activating similar innate immune responses as in infected cells (Chen and Hur, 2022; Stok et al., 2020). Accordingly, the innate immune and inflammatory response to dsRNAs underlie diverse pathologies from autoimmunity to neurodegeneration and to metabolic disorders (Saldi et al., 2019; Sud et al., 2016; Uggenti et al., 2019).

One family of innate immune receptors that shapes the cellular response to dsRNA are RIG-I-like receptors (RLRs) (Ablasser and Hur, 2020). RIG-I and MDA5, the two primary RLRs, are cytoplasmic proteins, are present in most vertebrate cells, and function as the first line of defense against a broad range of viruses. Upon dsRNA binding, RLRs multimerize and activate the signaling adaptor molecule MAVS by inducing MAVS multimerization (Hou et al., 2011; Peisley et al., 2011; Peisley et al., 2013; Zeng et al., 2010). Multimerized MAVS then triggers a cascade of biochemical events culminating in the activation of IRF3 and NF-κB, and subsequent induction of a largely group of antiviral genes, including type I interferons (IFNs).

In addition to transcriptional remodeling by the RLR pathway, foreign dsRNA also triggers other cellular changes, including assembly of molecular condensates known as stress granules (SGs) (Hofmann et al., 2021; Van Treeck and Parker, 2019). SG assembly is a highly conserved cellular phenomenon in eukaryotes, and is induced not only by dsRNA, but also by other cellular stress conditions, including heat shock and oxidative stress. These diverse stimuli activate several kinases, for example the dsRNA-dependent kinase PKR, which commonly phosphorylate the translational initiation factor eIF2α and suppress global protein synthesis to help cells recover from stress (Costa-Mattioli and Walter, 2020). SGs are formed when stalled ribosome-mRNA complexes accumulate and aggregate together with other cytoplasmic proteins, including the key nucleators G3BP1/2 and UBAP2L (Cirillo et al., 2020; Guillen-Boixet et al., 2020; Kedersha et al., 2016; Sanders et al., 2020; Yang et al., 2020). While SGs were initially thought to be the sites of translational suppression, more recent studies suggested that SG formation is not necessary for translational suppression (Bley et al., 2015; Mateju et al., 2020), raising questions about the physiological functions of the granule formation.

Multiple models have suggested a link between SGs and antiviral immune responses. One prominent theory is that SGs function as the signaling scaffold for RLRs (Oh et al., 2016; Onomoto et al., 2013). This is based on the observations that, upon infection, RLRs localized at SGs together with a subset of viral RNAs, and that knocking down G3BPs inhibited SG assembly and diminished induction of type I IFNs. This was in line with the previous reports that SGs are frequently targeted or altered by many viruses (Feng et al., 2014; McCormick and Khaperskyy, 2017; White and Lloyd, 2012), although often through indirect mechanisms such as limiting accessibility of viral dsRNA. However, other reports have raised questions whether SGs are in fact the sites of RLR activation. While a subset of viral RNAs were found to accumulate within SGs, dsRNAs appeared to be excluded (Langereis et al., 2013; Oh et al., 2016).

Additionally, SG-disrupting pharmacological agents (e.g. cycloheximide) did not impair RLR signaling, while other stressors, such as arsenite or heat-shock, triggered SGs and RLR colocalization without activating RLRs (Cadena et al., 2019; Langereis et al., 2013). Furthermore, SGs were proposed to suppress other innate immune pathways, such as NLRP3 inflammasome and MAPK signaling (Arimoto et al., 2008; Samir et al., 2019), which raises additional questions as to what role SGs play in innate immunity and whether there is a generalizable principle.

Here we report evidence supporting that SG’s primary function is to maintain cell homeostasis during infection. This function is mediated by two distinct mechanisms. First, SGs prevent excessive activation of RLR signaling and immune-mediated cell death. Second, SGs have cell-intrinsic activity that suppress viral replication in a manner independent of RLRs. These findings highlight the multi-functional nature of SGs in innate immunity.

## Results

### SG-deficient ΔG3BPs cells display hyperactivation of RLR signaling

In an effort to understand the role of SGs in RLR signaling, we first examined cellular response to *in vitro* transcribed 162 bp dsRNA harboring 5’-triphosphate groups (5’ppp), a known ligand that directly activates RIG-I (Cadena et al., 2019). Use of a viral dsRNA mimic ensures a potent stimulation of the RLR pathway in the absence of confounding factors such as viral antagonisms. We confirmed that SGs were formed in U2OS cells upon dsRNA transfection, as measured by immunofluorescence (IF) analysis of the SG markers G3BP1 or TIAR (Hofmann et al., 2021; Van Treeck and Parker, 2019) (Figures 1A and S1A-B). We also confirmed that SGs recruited RLRs (RIG-I and MDA5), their signaling adaptor MAVS and downstream signaling molecules, including TRAF proteins and TBK1 (Figures 1A and S1B). However, SG localization of dsRNA was minimal regardless of whether dsRNA was delivered to the cytoplasm by cationic lipid transfection or by electroporation (Figure 1A, bottom). The lack of dsRNA colocalization is not in line with the notion that SGs are the sites of RLR activation.

**Figure 1.**
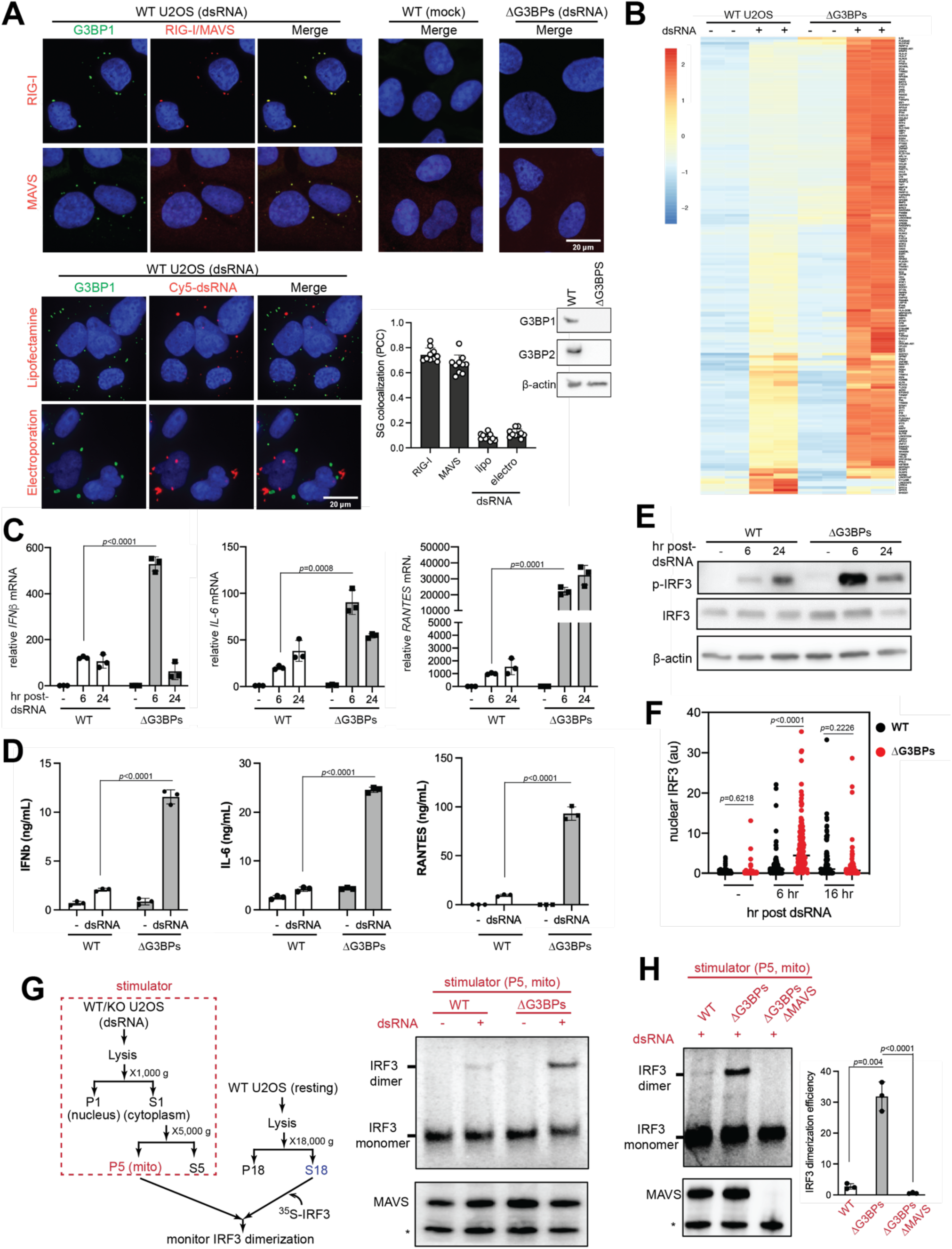
RLR signaling is hyperactive in SG-deficient ΔG3BPs cells. **A.** Immunofluorescence (IF) analysis of RIG-I, MAVS and dsRNA (red) with G3BP1 (green) in U2OS cells (WT vs ΔG3BPs). See Figure S1A for antibody validation. Cells were transfected with 162 bp dsRNA containing 5’ppp (500 ng/ml) for 6 hrs prior to imaging. For dsRNA imaging, 162 bp dsRNA 3’-labeled with Cy5 was introduced into cells by either lipofectamine transfection or electroporation (see Figure S1C for RLR signaling in response to Cy5-dsRNA). Unless mentioned otherwise, unlabeled dsRNA was introduced to cells by lipofectamine transfection throughout the manuscript. Cell nuclei were stained with Hoechst 3342. Bottom right: SG colocalization was measured by Pearson colocalization coefficient (PCC) between G3BP1 foci and indicated molecules from 10 fields of view. **B.** Heatmap of z-scores displaying differentially expressed genes in WT vs ΔG3BPs U2OS cells. Cells were transfected with 162 bp dsRNA with 5’ppp (500 ng/ml) for 6 hrs. Genes showing log2-fold change (lfc2) >2 (with *p*_adj<0.05) upon dsRNA stimulation in a MAVS-dependent manner were shown. The list of MAVS-dependent genes was compiled from the comparison of RNA-seq data from ΔG3BPs and ΔG3BPsΔMAVS (Figure S2B). All genes were shown in Figure S2A. **C.** Levels of *IFNβ* (left), *IL-6* (middle), and *RANTES* (right) mRNAs. U2OS cells were transfected with dsRNA as in (B) and were analyzed 6 or 24 hr post-dsRNA. **D.** Levels of secreted IFNβ (left), IL-6 (middle), and RANTES (right) as measured by ELISA. U2OS cells were transfected with dsRNA as in (B) and were analyzed at 6 hr post-dsRNA. **E.** Activation state of IRF3, as measured by its phosphorylation level. U2OS cells were stimulated as in (B). **F.** Activation state of IRF3, as measured by its nuclear translocation. U2OS cells were stained with anti-IRF3 antibody at indicated timepoints and the level of nuclear IRF3 signal was quantitated (a.u. indicates arbitrary unit). Each data point represents a nucleus (n=61-179). DAPI staining was used for defining nuclear boundary. Of note, the latest time-point for this experiment was 16 hr, rather than 24 hr to avoid pronounced cell detachment due to cell death (to be discussed in Figure 4). **G.** Activation state of MAVS, as measured by cell-free IRF3 dimerization assay. Mitochondrial fraction (P5) containing MAVS was isolated from U2OS cells stimulated with dsRNA as in (B), and then mixed with a common pool of cytosolic extract (S18) from unstimulated WT U2OS cells and in vitro translated ^35^S-IRF3. Dimerization of ^35^S-IRF3 was analyzed by native gel assay. The level of MAVS was analyzed to ensure that an equivalent amount of MAVS was used in all samples. * indicates mini-MAVS. **H.** Cell-free IRF3 dimerization assay, comparing the activity of the mitochondrial fraction isolated from WT, ΔG3BPs and ΔG3BPs/ΔMAVS U2OS cells. Right: relative level of the dimeric IRF3. Data are presented in means ± SD. *p* values were calculated using two-tailed unpaired Student’s t test (ns, *p*>0.05). RNA-seq results contain 2 biological repeats and were confirmed by two independent experiments. All other data are representative of at least three independent experiments.). Raw data for the heatmap can be found in the supplemental file (data 1).

We next compared RLR signaling in wild-type (WT) and SG-deficient cells by examining their transcriptome at two time points (6 and 24 hr post-dsRNA). To this end, we used cells deficient in the two key nucleators G3BP1 and 2 (ΔG3BPs) (Guillen-Boixet et al., 2020; Kedersha et al., 2016; Sanders et al., 2020; Yang et al., 2020). ΔG3BPs cells were deficient in formation of SGs and RIG-I/MAVS foci upon dsRNA introduction (Figure 1A). Remarkably, ΔG3BPs cells displayed enhanced antiviral signaling than WT cells at 6 hr post-dsRNA transfection (Figures 1B and S2A). This was confirmed by independent analysis of mRNA levels of select few cytokines (IFNβ, IL-6 and RANTES, Figure 1C) and their secreted protein levels (Figure 1D). G3BPs complementation in ΔG3BPs cells restored SG formation and suppressed RLR signaling (Figure S2C), further supporting the role of G3BPs in suppressing RLR signaling.

In contrast to the immediate response to dsRNA, RLR signaling at 24 hr post-dsRNA showed more complex, gene-specific patterns in ΔG3BPs cells. While most dsRNA-induced genes increased in expression from 6 to 24 hr post-dsRNA, a subset of genes, including *IFNβ*, markedly decreased from 6 hr to 24 hr (Figure S2D). Similarly, RT-qPCR measurement showed that the spike in the *IFNβ* mRNA level in ΔG3BPs cells at 6 hr was followed by a sharp decline at 24 hr to the level comparable to the WT level (Figure 1C). In contrast, *RANTES* and *IL-6* mRNAs remained higher in ΔG3BPs cells at both 6 and 24 hr (Figure 1C).

Given the complex behavior of gene induction, we next examined RLR signaling by measuring the activation state of the upstream signaling molecules IRF3 and MAVS. IRF3 is the transcription factor responsible for *IFNβ* induction and its activation requires IRF3 phosphorylation, dimerization and nuclear translocation. Analysis of the levels of p-IRF3 and nuclear IRF3 showed that IRF3 was more active in ΔG3BPs than in WT cells at 6 hr post-dsRNA (Figures 1E & 1F). However, at 24 hr post-dsRNA, we observed a sharp drop in the levels of p-IRF3 and nuclear IRF3 in ΔG3BPs cells, mirroring the pattern of *IFNβ* mRNA induction. As will be discussed in Figure 4, this decline of the IRF3 activity was due to negative feedback regulation and cell death that was enhanced in ΔG3BPs cells as a result of increased RLR signaling at early time points.

We next examined the activation state of MAVS using a previously established cell free assay (Hou et al., 2011; Zeng et al., 2010). The mitochondrial fraction (P5) containing MAVS was extracted from dsRNA-stimulated WT or ΔG3BPs cells, and the signaling potential of MAVS was measured by incubating P5 with cytosolic fraction (S18) from unstimulated WT cells, which provided a common pool of downstream signaling molecules in the resting state (Figure 1G, left). In vitro-translated, inactive ^35^S-labeled IRF3 was added to the mixture in order to measure MAVS’ ability to activate IRF3, as visualized by the monomer-to-dimer transition of ^35^S-IRF3 in the native gel. We found that only P5 from dsRNA-stimulated cells could activate ^35^S-IRF3 and that P5 from ΔG3BPs cells was more potent than WT P5 (Figure 1G, right). ^35^S-IRF3 activation was mediated entirely by MAVS in P5, since P5 from ΔG3BPsΔMAVS was unable to stimulate ^35^S-IRF3 (Figure 1H). Altogether, these results showed that the RLR signaling pathway is more potently activated by dsRNA in ΔG3BPs than in WT cells, as measured by RLR-induced cytokine levels, global transcriptome, and activation states of IRF3 and MAVS.

We next asked whether hyper-activation of RLR signaling in ΔG3BPs cells at an early timepoint is generalizable to other cell types and whether this is independent of the method of dsRNA delivery or the type of dsRNA. As with U2OS cells, A549 and HeLa cells also formed SGs upon dsRNA introduction (Figure S3A). These cells also displayed hyper-activation of RLR signaling in ΔG3BPs than in WT cells 4 hrs following dsRNA introduction (Figure S3B & S3C). Comparing different methods of dsRNA delivery, we found that ΔG3BPs cells consistently showed higher levels of RLR signaling either by electroporation (Figure S3D) or lipofectamine transfection of dsRNA (Figure 1). Poly I:C, a synthetic dsRNA mimetic, also triggered more potent RLR signaling in ΔG3BPs cells (Figure S3E). These data could be further expanded to viral infections (to be discussed in Figures 6 and S6). However, antiviral signaling in response to cGAMP, which activates the cGAS-STING pathway without SGs, was comparable in WT and ΔG3BPs cells (Figure S3F). These results show that G3BPs suppress RLR signaling in response to dsRNA stimulation in multiple cell lines, but this is not due to a generic decrease in antiviral signaling efficiency.

### SGs suppress RLR signaling

To examine whether the observed effect of G3BPs on RLR signaling is indeed mediated by SGs or other potential functions of G3BPs, we next examined two other genetic models for SG deficiency, ΔUBAP2L and ΔPKR. Like G3BPs, UBAP2L is an essential nucleator for SGs, the deficiency of which impairs SG formation (Cirillo et al., 2020; Sanders et al., 2020). In contrast to G3BPs and UBAP2L, PKR is not directly involved in SG assembly, but triggers translational suppression in response to dsRNA, a pre-requisite for SG nucleation (Hofmann et al., 2021; Van Treeck and Parker, 2019). Previous studies suggested that certain granules distinct from SGs can form in ΔPKR cells upon dsRNA stimulation (Burke et al., 2020). In keeping with this report, we found that ΔPKR cells still form G3BP1 foci, but they did not show enrichment of MAVS or the other SG marker TIAR (Figure 2A) and were significantly smaller in size (Figure 2B, top).

**Figure 2.**
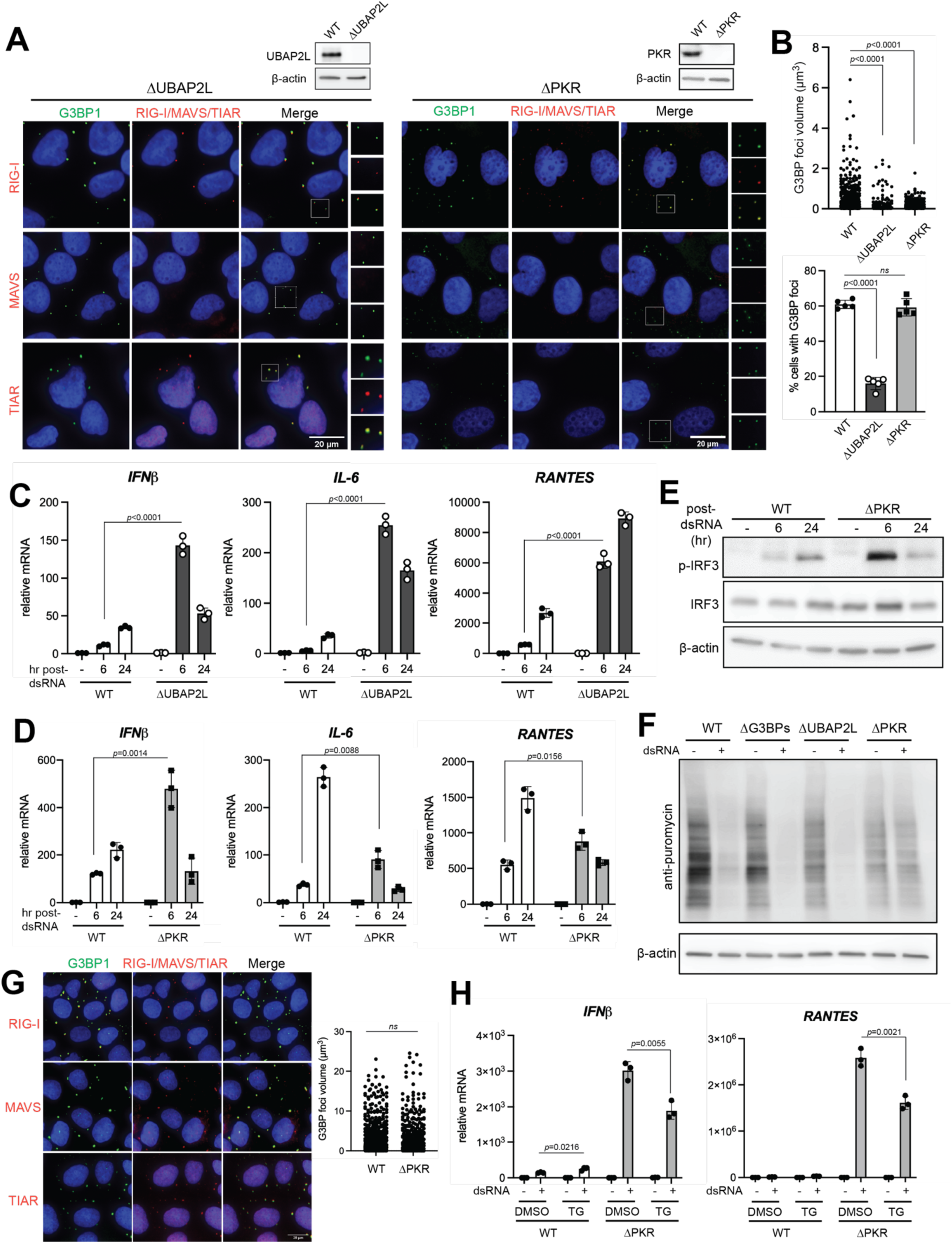
SG deficiency leads to hyperactivation of RLR signaling. **A.** Immunofluorescence analysis of RIG-I, MAVS, TIAR (red) and G3BP1 (green) in ΔUBAP2L and ΔPKR U2OS cells. Cells were transfected with 162 bp dsRNA at 500 ng/ml for 6 hrs. Insets show G3BP1 foci at higher magnifications. **B.** G3BP1 foci size and frequency in ΔUBAP2L and ΔPKR U2OS cells. Foci size was quantitated for at least 200 randomly selected granules from Z-stack images (0.15 μm step size). Foci frequency was measured from 5 fields of view. **C.** Antiviral signaling in U2OS cells (WT vs ΔUBAP2L) in response to dsRNA transfection (500 ng/ml), as measured by the level of *IFNβ* (left), *IL-6* (center) and *RANTES* (right) mRNAs at 6 or 24 hr post-dsRNA. **D.** Same as (C) comparing WT and ΔPKR U2OS cells. **E.** IRF3 phosphorylation in U2OS cells (WT vs ΔPKR). Cells were stimulated as in (D). **F.** Level of protein synthesis as measured by puromycin incorporation (SUnSET assay). U2OS cells were transfected with dsRNA (500 ng/ml) for 6 hrs and pulsed with puromycin (1 μg/ml) for 15 mins prior to analysis of nascent peptide using anti-puromycin WB. **G.** Colocalization of RIG-I, MAVS and TIAR (red) with G3BP1 (green) in U2OSΔPKR cells upon treatment with TG (1 μM) without dsRNA. G3BP1 foci size was quantitated for at least 600 randomly selected granules from Z-stack images (0.15 μm step size). **H.** Antiviral signaling in U2OS cells (WT vs ΔPKR) in response to dsRNA (500 ng/ml), in the presence and absence of TG. Cells were treated with TG (1 μM) at 1 hr post-dsRNA and harvested 6 hr post-dsRNA for *IFNβ* and *RANTES* mRNA measurement. Data are presented in means ± SD. *p* values were calculated using two-tailed unpaired Student’s t test (ns, *p*>0.05). Data are representative of three independent experiments.

ΔUBAP2L cells also displayed G3BP1 foci upon dsRNA stimulation, but these foci lacked MAVS (Figure 2A) and were smaller in size and less frequent (Figure 2B). Thus, both ΔPKR and ΔUBAP2L cells can form G3BP1 foci but they are distinct from conventional SGs in WT cells. Importantly, both ΔUBAP2L and ΔPKR cells showed hyperactivation of RLR signaling at 6 hr post-dsRNA (Figures 2C & 2D), mirroring that of ΔG3BPs cells. Measurement of p-IRF3 also confirmed hyperactivation of the RLR pathway in ΔPKR cells at 6 hr (Figure 2E). Thus, analysis of three distinct SG-deficient backgrounds (ΔG3BPs, ΔUBAP2L and ΔPKR) commonly suggest that SGs suppress RLR signaling, at least at early time points.

At 24 hr post-dsRNA, however, ΔPKR cells diverged from ΔUBAP2L and ΔG3BPs cells in gene induction patterns, in particular those of *IL-6* and *RANTES*. Both ΔUBAP2L and ΔG3BP cells showed sustained activation of *IL-6* and *RANTES,* displaying higher levels than WT cells at both 6 and 24 hr. ΔPKR cells, on the other hand, showed dramatic reduction in both genes from 6 to 24 hr, dropping below the WT levels (Figures 2C & 2D). We suspect that this divergence reflects the fact that ΔPKR affects both translation and SG formation, while ΔG3BPs and ΔUBAP2L selectively affect SG formation without altering translational control (see Figure 2F for comparison of translational activity using SUnSET (Schmidt et al., 2009)). Since inhibiting translation alone (without SG induction) has a positive effect on innate immune signaling (Dalet et al., 2017; Deng et al., 2004; McAllister et al., 2012), lack of translational inhibition in ΔPKR may be responsible for reversing the positive effect of the SG deficiency at later time point.

We next asked whether the RLR-suppressive function is specific to dsRNA-triggered SGs or whether SGs triggered by other types of stress, such as ER stress, can also repress RLR signaling. One of the best characterized ER stressors is thapsigargin (TG), which triggers SGs by activating PERK, instead of PKR. Accordingly, TG stimulation triggers SGs in ΔPKR cells, which colocalize with RIG-I, MAVS and TIAR foci (Figure 2G). Since ΔPKR does not form SGs in response to dsRNA, ΔPKR cells offered a unique opportunity to test the effect of TG-triggered SGs on RLR signaling. TG treatment in ΔPKR cell reduced RLR signaling at 6 hr post-dsRNA (Figure 2H). Similar suppression was not observed in WT cells where dsRNA alone triggered SGs (Figure 2H). Altogether, these results suggest that SGs suppress RLR signaling, regardless of whether SGs are formed by PKR- or PERK-mediated pathway.

### SGs suppress other dsRNA-dependent innate immune pathways

We next asked whether SGs can also affect other innate immune pathways, such as those mediated by two other families of dsRNA sensors, PKR and OAS enzymes (Figure 3A). As discussed earlier, PKR activation by dsRNA leads to global suppression of protein synthesis through phosphorylation of eIF2α (Pfeller et al., 2011). Activation of OASes by dsRNA also leads to restriction of protein synthesis, but through a different downstream effector, RNase L – a highly potent and non-specific ribonuclease that cleaves rRNAs, tRNAs and mRNAs (Kristiansen et al., 2011). Together with RLRs, PKR and OAS pathways contribute to antiviral innate immunity, restrict cell proliferation and cause cell death in extreme cases (Banerjee et al., 2019; Gannon et al., 2018; Ishizuka et al., 2019; Li et al., 2017). Interestingly, we found that PKR, OAS3 and RNase L are all highly enriched within SGs (Figures 3B-C), in line with previous reports (Onomoto et al., 2012; Reineke and Lloyd, 2015). This prompted us to ask whether their activities are also regulated by SGs, possibly through molecular sequestration.

**Figure 3.**
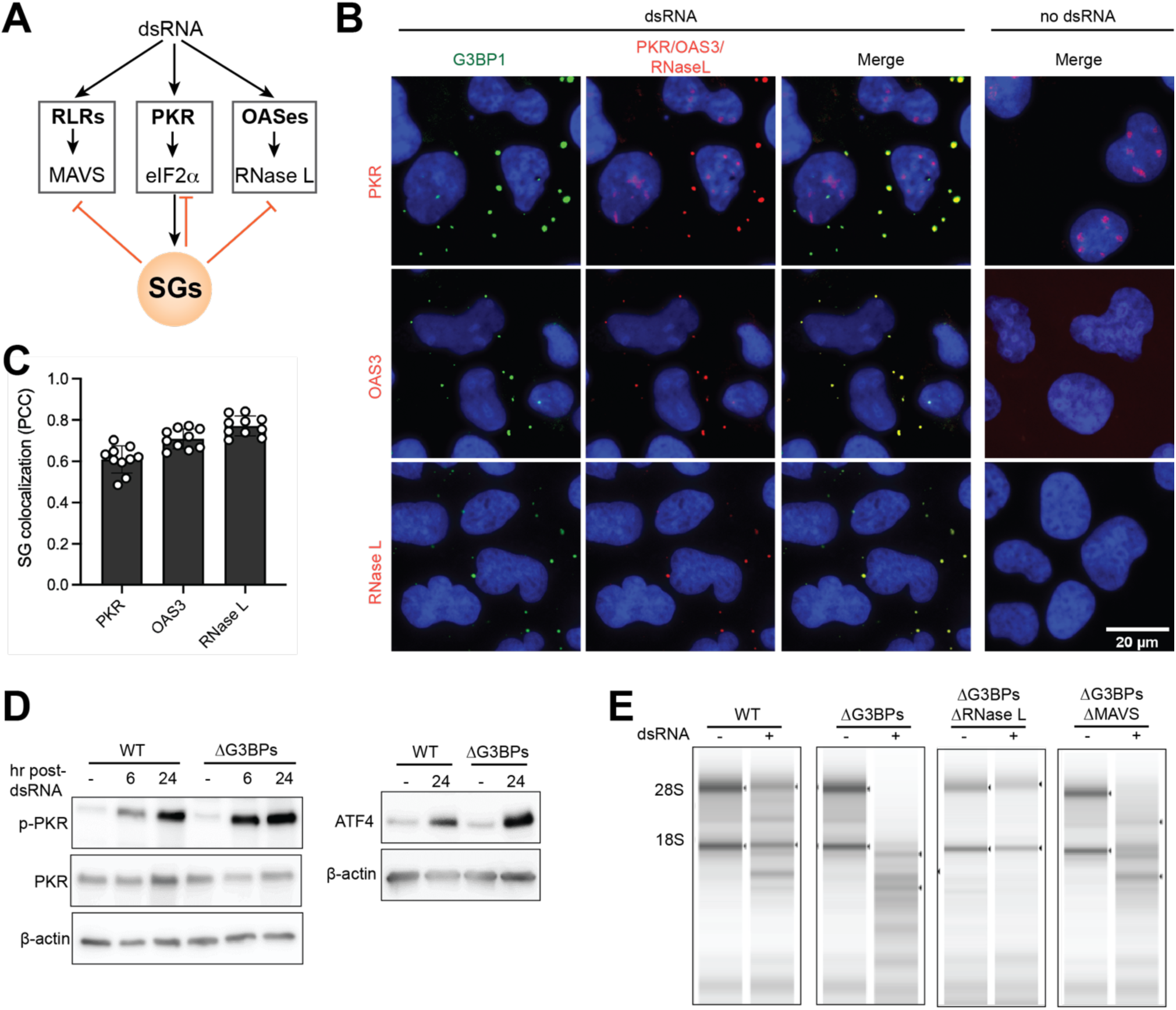
SGs also suppress PKR and OAS pathways. **A.** Schematic of dsRNA-dependent innate immune pathways. Besides RLRs, most vertebrate cells express two other dsRNA sensors, PKR and OASes. PKR activation leads to eIF2α-mediated translational suppression and subsequent formation of SG. OASes activation, on the other hand, leads to RNase L-mediated degradation of bulk cytosolic RNAs (including mRNA, rRNA and tRNA). Our results in Figures 1-3 suggest that SGs suppress all three dsRNA-dependent innate immune pathways. **B.** Immunofluorescence analysis of PKR, OAS3, RNase L (red) and G3BP1 (green) in U2OS cells. Cells were transfected with 162 bp dsRNA (500 ng/ml) for 6 hrs prior to imaging. **C.** SG colocalization was measured by Pearson colocalization coefficient (PCC) between G3BP1 foci and indicated molecules from 10 fields of view. **D.** PKR activity in WT vs. ΔG3BPs U2OS cells as measured by PKR phosphorylation (left) and ATF4 expression (right) at indicated time points. **E.** RNase L activity in WT vs. ΔG3BPs U2OS cells as measured by rRNA degradation. Total RNA was isolated 24 hr post-dsRNA and was analyzed by Agilent TapeStation.

**Figure 4.**
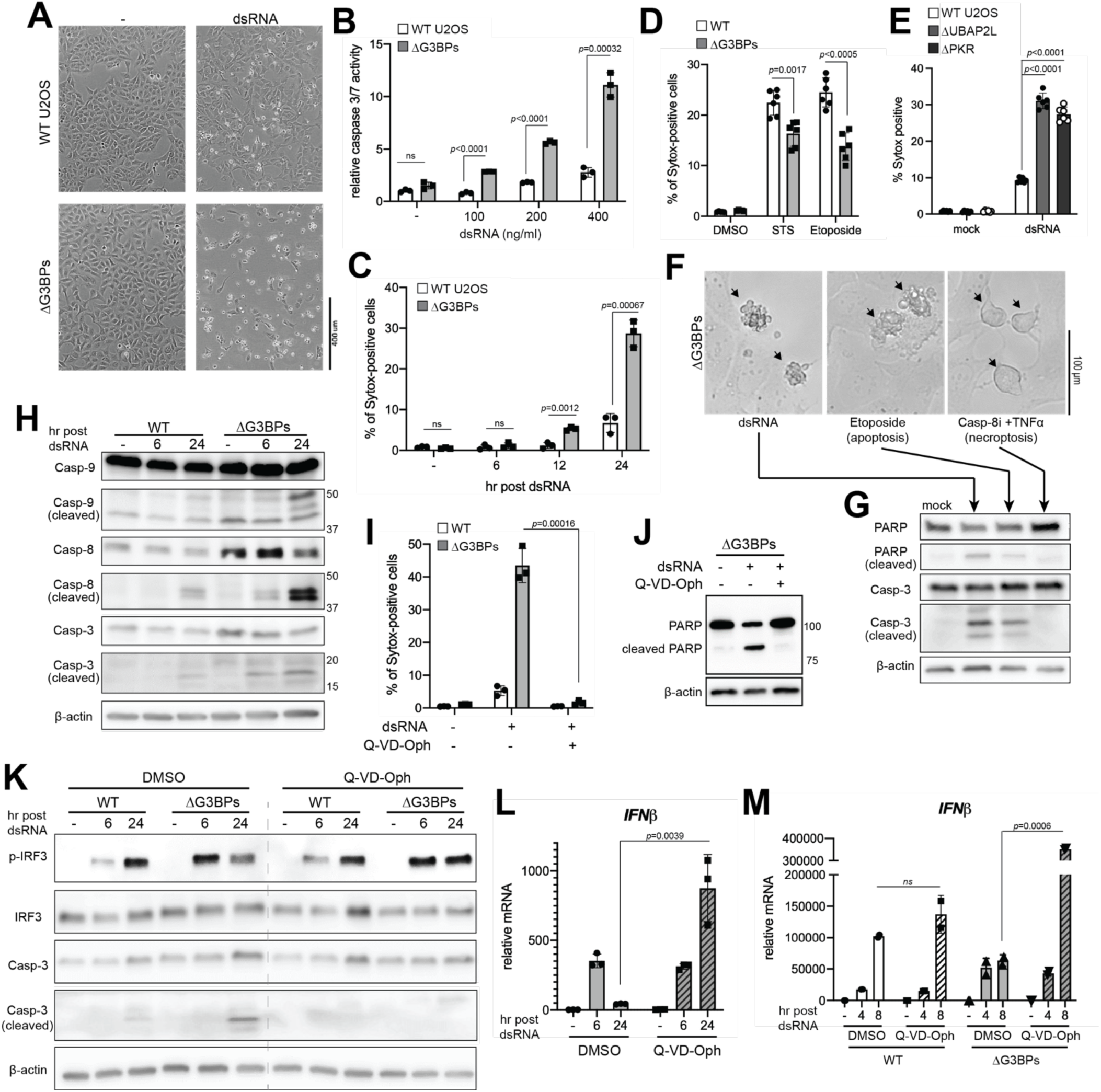
SGs dampen dsRNA-triggered apoptosis and the consequent negative feedback regulation of IRF3. **A.** Cell death in WT vs. ΔG3BPs U2OS cells as examined by bright-field microscopy. Cells were transfected with 162 bp dsRNA (500 ng/ml) as in Figure 1B and analyzed at 24 hr post-dsRNA. **B.** Cell death as measured by Caspase-3/7 activity. U2OS cells were transfected with an increasing concentration of 162 bp dsRNA and analyzed at 24 hr post-dsRNA. **C.** Cell death as measured by Sytox uptake at indicated time points. Percentage of Sytox-positive cells among Hoechst 3342-positive cells was plotted. **D.** Cell death in response to staurosporin (STS) and epotoside, as measured by Sytox uptake. U2OS cells were treated with STS (1 μM) or etoposide (20 μM) for 24 hrs before Sytox analysis. **E.** Cell death in WT, ΔUBAP2L and ΔPKR U2OS cells, as measured by Sytox uptake. Cells were transfected with 162 bp dsRNA (500 ng/ml) and were analyzed at 24 hr post-dsRNA. **F.** Comparison of cell death triggered by dsRNA, etoposide and a combination of caspase-8 inhibitor (Z-IETD-FMK, Casp-8i) and TNFα. Etoposide was used as a known trigger for apoptosis, while Casp-8i+TNFα was for necroptosis. **G.** Analysis of PARP and caspase-3 (Casp-3) cleavage using samples from (F). **H.** Apoptotic caspase cleavage in WT vs ΔG3BPs U2OS cells at 6 or 24 hr post-dsRNA. **I.** Effect of pan-caspase inhibitor (Q-VD-OPh) on dsRNA-triggered cell death, as measured by Sytox uptake at 24 hr post-dsRNA. U2OS cells were treated with Q-VD-OPh (10 μM) 1 hr pre-dsRNA. **J.** Effect of pan-caspase inhibitor (Q-VD-OPh, 10 μM) on dsRNA-triggered cell death, as measured by PARP cleavage in U2OSΔG3BPs cells at 24 hr post-dsRNA. **K.** Effect of Q-VD-OPh on IRF3 phosphorylation and caspase-3 cleavage. **L.** Effect of Q-VD-OPh on *IFNβ* mRNA induction in U2OSΔG3BPs cells. **M.** Effect of Q-VD-OPh on *IFNβ* mRNA induction in A549 cells. Cells were transfected with 162 bp dsRNA (500 ng/ml) and treated with Q-VD-OPh (10 μM) 1 hr pre-dsRNA. Data are presented in means ± SD. *p* values were calculated using two-tailed unpaired Student’s t test (ns, *p*>0.05). All data are representative of three independent experiments.

We first examined the PKR activity as measured by the levels of p-PKR and ATF4, which were known to increase upon PKR activation (Costa-Mattioli and Walter, 2020), and found more potent activation of PKR in ΔG3BPs cells than in WT cells (Figure 3D). Given that PKR is an upstream inducer of SGs, this result suggests that SG is a negative feedback regulator of PKR. To measure the OAS-RNase L pathway activity, we examined the integrity of total cellular RNA, most prominently 18S and 28S rRNAs. We found that RNase L is more active in ΔG3BPs cells than in WT cells to the extent that nearly all rRNAs were degraded in ΔG3BPs cells by 24 hr post-dsRNA (Figure 3E). Since OASes were known to be induced by type I IFNs, and thus RLR signaling, we examined whether enhanced RLR signaling in ΔG3BPs could explain the hyperactivity of RNase L. Knocking out MAVS only partly rescued RNA integrity of ΔG3BPs cells (Figure 3E), suggesting that SG-deficiency causes enhanced activity of the OAS-RNase L pathway largely independent of RLRs. Altogether, these results suggest that SGs inhibit all three dsRNA-dependent innate immune pathways involving RLRs, PKR and OASes (Figure 3A).

### Lack of SG leads to apoptosis and the consequent suppression of IRF3 at later time points

In addition to hyperactivation of innate immune signaling pathways, we also found that ΔG3BPs cells underwent pronounced cell death upon dsRNA stimulation, as measured by the loss of cell attachment to the surface (Figure 4A). This was further confirmed by Sytox uptake, caspase-3/7 activity assay and LIVE/DEAD dye staining (Figures 4B-4C and S4A). While high dose of dsRNA is known to trigger cell death, ΔG3BPs cells underwent significantly more pronounced cell death than WT cells at all doses of dsRNA tested (Figure 4B). This dsRNA-triggered cell death progressed gradually over a 24-hr time course (Figure 4C). The hypersensitivity of ΔG3BPs cell viability is dependent on the cell death trigger, as other stimuli, such as etoposide and staurosporine, did not trigger greater cell death in ΔG3BPs than WT cells (Figure 4D). Similar increase in dsRNA-triggered cell death was observed with A549ΔG3BPs (Figures S4B-D) and HeLaΔG3BPs cells (Figures S4E-G). Moreover, ΔUBAP2L and ΔPKR cells also underwent increased cell death upon dsRNA stimulation (Figure 4E), further supporting the notion that the lack of SGs is responsible for the hypersensitivity to dsRNA.

Detailed analysis of dsRNA-induced cell death in ΔG3BPs showed morphologic features similar to apoptotic blebs (Figure 4F, left and center) but distinct from necroptotic cells (Figure 4F, right). We also observed cleavage of PARP and caspase-3, which again mirrored that of apoptotic, not necroptotic cells (Figure 4G). Stimulation of ΔG3BPs cells with dsRNA triggered cleavage of multiple caspases, including caspase-3, -8, -9 (Figure 4H), but not caspase-1 (Figure S4H). Consistent with the notion that ΔG3BPs cells undergo apoptosis, a pan-caspase inhibitor (Q-VD-Oph) completely blocked dsRNA-dependent cell death and cleavage of PARP in ΔG3BPs (Figures 4I & 4J), but a pyroptosis inhibitor (disulfiram) or necroptosis inhibitor (MLKL inhibitor) did not (Figure S4I). Thus, dsRNA-mediated cell death in ΔG3BPs is caspase-dependent apoptosis.

We next asked whether apoptosis or caspase activation was the reason for the marked decline in the IRF3 activity in ΔG3BPs cells at later time points (Figures 1C and 1E). This hypothesis was based on previous reports showing that caspases can suppress IRF3 signaling by cleaving MAVS, IRF3 and/or RIP1 (Ning et al., 2019; Rajput et al., 2011; Rongvaux et al., 2014). While we did not observe clear signs of caspase-dependent cleavage of RIG-I, MAVS and IRF3 (Figure S4J), Q-VD-Oph significantly improved the level of p-IRF3 at 24 hr without significantly altering it at 6 hr (Figure 4K). This was in line with the observation that caspase activity was high at 24 hr but was minimal at 6 hr (Figure 4H). Similar effect of Q-VD-Oph was observed on the level of *IFNβ* mRNA in U2OSΔG3BPs (Figure 4L) and A549ΔG3BPs cells (Figure 4M). In keeping with minimal activation of caspases in WT U2OS cells, Q-VD-Oph did not affect RLR signaling in WT cells (Figures 4K and 4M). Collectively, these data suggest that the lack of SGs results in hyperactivation of caspases in response to dsRNA, which causes a marked decline in the IRF3 activity at later time points. In other words, SGs prevent dsRNA-induced caspase activation and cell death, which allows cells to minimize caspase-dependent regulation of RLR-MAVS-IRF signaling and enables sustained immune signaling.

### SGs minimize cell death by preventing overstimulation of innate immune pathways by dsRNA

We next investigated why SG deficiency leads to increased cell death in response to dsRNA stimulation. We first examined the potential role of the RLR-MAVS pathway, as it has previously been shown to contribute to dsRNA-dependent cell death (Besch et al., 2009; Lei et al., 2009). Knocking out MAVS largely rescued cell viability in U2OSΔG3BPs (Figure 5A) and A549ΔG3BPs (Figure S5A) upon dsRNA stimulation. Knocking out MAVS also significantly reduced cleavage of caspase-3, -8 and -9, albeit not to completion (Figure 5B). To examine whether the IRF3-IFNα/β signaling axis plays a role in MAVS-mediated cell death, we knocked out IRF3 in U2OSΔG3BPs cells, which completely abrogated *IFNβ* induction (Figure 5C, left). However, knocking out IRF3 did not relieve dsRNA-triggered cell death in ΔG3BPs cells (Figure 5A). In addition, inhibitors of TBK1 and JAK that respectively block IRF3 activation and the downstream actions of type I IFNs did not alleviate dsRNA-triggered cell death in ΔG3BPs (Figure S5B). These results suggest that the cell death in ΔG3BPs is mediated by a MAVS-dependent, but IRF3- and IFN-independent mechanism.

**Figure 5.**
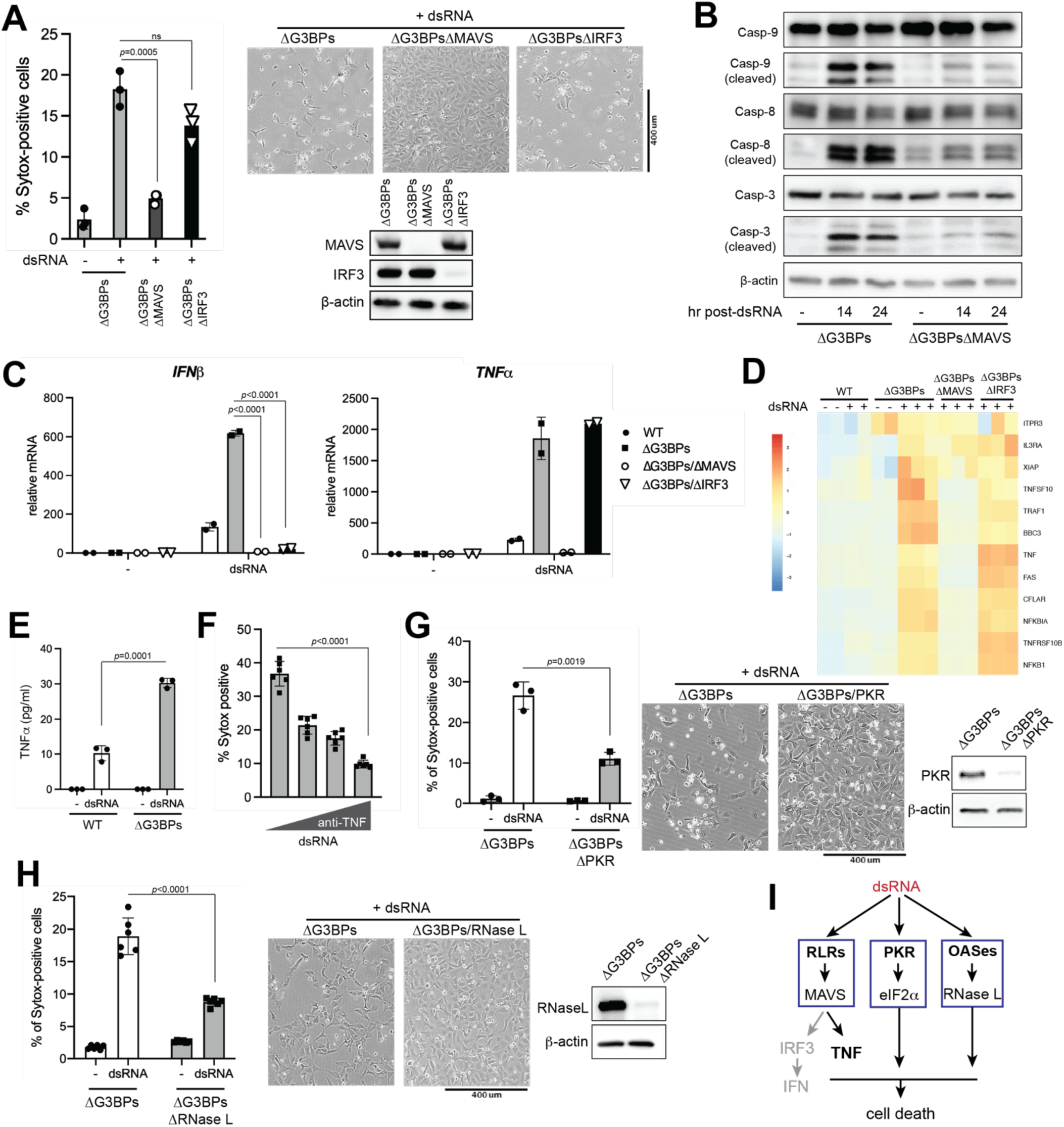
SGs prevent dsRNA-triggered cell death by suppressing RLR, PKR and OAS pathways. **A.** Cell death in WT, ΔG3BPs, ΔG3BPsΔMAVS and ΔG3BPsΔIRF3 U2OS as measured by Sytox uptake (left) and bright field microscopy (right) at 24 hr post-dsRNA. Cells were transfected with 162 bp dsRNA as in Figure 1B. **B.** Apoptotic caspase cleavage in ΔG3BPs and ΔG3BPsΔMAVS U2OS cells at 14 or 24 hr post-dsRNA. **C.** Levels of *IFNβ* and *TNFα* mRNAs in U2OS cells at 6 hr post-dsRNA. **D.** Heat map of z-scores for differentially expressed genes in apoptosis pathway (KEGG pathway hsa04210) in U2OS cells at 6 hr post-dsRNA stimulation. **E.** Level of secreted TNFα in U2OS cells 6 hr post-dsRNA. **F.** Effect of anti-TNFα antibody on dsRNA-triggered cell death in U2OSΔG3BPs. Cells were pre-treated with anti-TNFα antibody (0.01, 0.1 and 1 μg/ml) 30 min prior to transfection with dsRNA. Cell death was measured by Sytox uptake at 24 hr post-dsRNA. **G.** Cell death in ΔG3BPs and ΔG3BPsΔPKR as measured by Sytox uptake (left) and bright field microscopy (right) at 24 hr post-dsRNA. **H.** Cell death in ΔG3BPs and ΔG3BPsΔRNase L as measured by Sytox uptake (left) and bright field microscopy (right) at 24 hr post-dsRNA stimulation. **I.** Schematic for dsRNA-induced cell death in ΔG3BPs cells. The lack of SGs make ΔG3BPs cells hypersensitive to dsRNA, resulting in more potent activation of RLR, PKR and OASes pathways upon dsRNA stimulation. The TNFα signaling branch (but not the IRF3-IFN branch) in the RLR-MAVS pathway makes the primary contribution to cell death in U2OS cells. The PKR and OASes-RNase L pathways also contribute to cell death, likely through their roles in suppressing global protein synthesis. Data are presented in means ± SD. *p* values were calculated using two-tailed unpaired Student’s t test (ns, *p*>0.05). All data are representative of three independent experiments.

**Figure 6.**
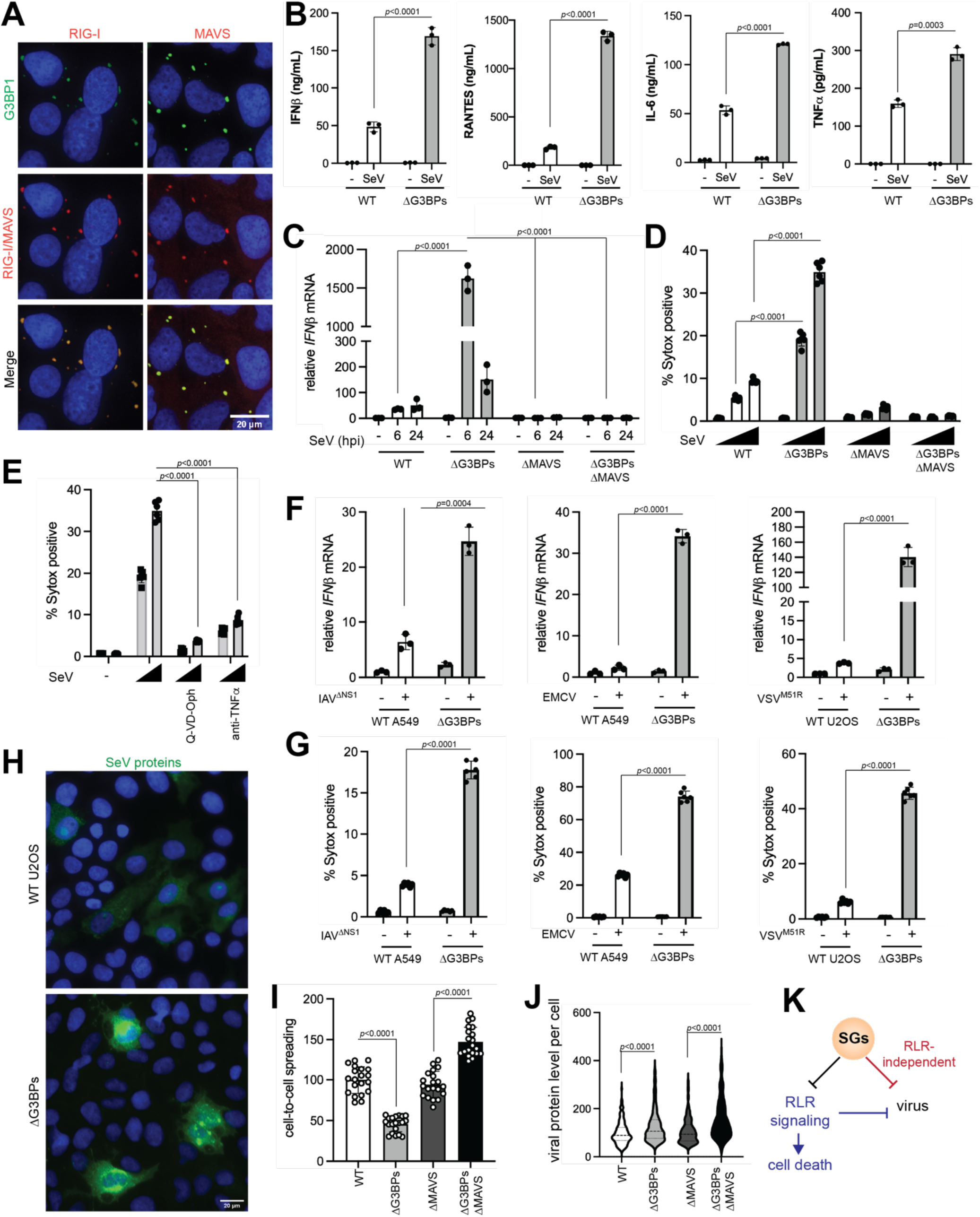
SGs suppress RLR signaling during viral infection, while restricting viral replication independent of RLRs. A. Immunofluorescence (IF) analysis of RIG-I, MAVS (red) and G3BP1 (green) in U2OS cells. Cells were infected with SeV (100 HA/ml) for 20 hrs prior to imaging. B. Levels of secreted IFNβ, IL-6, RANTES and TNFα as measured by ELISA. U2OS cells were infected with SeV (100 HA/ml) and were analyzed 6 hr post-infection (hpi). C. Antiviral signaling in U2OS cells during SeV infection (MOI=1.0), as measured by *IFNβ* mRNA at 6 or 24 hpi. D. Cell death in U2OS cells during SeV infection (MOI=0, 0.1, and 1.0), as measured by Sytox uptake at 24 hpi. E. Effect of anti-TNF and Q-VD-Oph on cell death in U2OSΔG3BPs cells upon SeV infection. Cells were infected with SeV (MOI=0.1, and 1.0), treated with inhibitors 1 hpi and analyzed at 24 hpi. F. Antiviral signaling upon infection with IAV^ΔNS1^, EMCV and VSV^M51R^, as measured by *IFNβ* mRNA. A549 cells were infected with IAV^ΔNS1^ (MOI=0.1) and EMCV (MOI=0.1), whereas U2OS cells were infected with VSV^M51R^ (MOI=1). Cells were harvested at 24 hpi for IAV^ΔNS1^ and 6 hpi for EMCV and VSV^M51R^. See Figure S6 for more comprehensive analysis with different MOIs and time of analysis. Note that IAV^ΔNS1^ infection was analyzed in both A549 and U2OS cells, which showed consistent results. G. Cell death upon infection with IAV^ΔNS1^, EMCV and VSV^M51R^, as measured by Sytox uptake. Cells were infected with viruses as in (F) and analyzed at 24 hpi. H. Immunofluorescent (IF) images of SeV proteins (green) in WT vs. ΔG3BPs U2OS cells. U2OS cells were infected with SeV (MOI=1) and stained with anti-SeV serum at 18 hpi. I. Relative cell-to-cell spreading of SeV. Cells were infected and number of cells above the background fluorescence (i.e. mock infection staining) per field of view were analyzed. Each data point represents a field of view (n=20). Data were normalized against the WT average value. J. Relative level of SeV protein staining in infected cells (MOI=1). Corrected total cell fluorescence (CTCF) was measured at 18 hpi. Each data point represents infected cell (n=200). Data were normalized against the WT average value. K. Schematic summarizing the dual function of SGs in (i) suppressing RLR signaling and (ii) restricting viral replication independent of the RLR pathway. Both functions converge on maintaining cell homeostasis. Data are presented in means ± SD. All data are representative of at least three independent experiments. *p* values were calculated using two-tailed unpaired Student’s t test (ns, *p*>0.05).

To identify the potential apoptosis-triggering factors downstream of MAVS, we looked for apoptosis-related genes among those up-regulated by dsRNA through the RLR pathway (Figure 5D). Several pro-apoptotic genes, including TNF (TNFα), FAS and TNFRSF10B (DR5), were hyperinduced in ΔG3BPs than in WT cells, and their induction was dependent on MAVS but not IRF3 (Figures 5C and 5D). Consistent with this, we detected markedly elevated secretion of TNFα in ΔG3BPs cells compared to WT cells (Figure 5E). Blocking TNFα signaling using anti-TNFα significantly relieved cell death in ΔG3BPs upon dsRNA stimulation (Figure 5F). However, treatment with TNFα alone did not induce cell death (Figure S5C), suggesting that TNFα cooperates with other factors to induce apoptosis in ΔG3BPs cells.

Given that knocking out MAVS did not completely rescue cell viability, we next examined the potential role of PKR and OAS-RNase L pathways, which were also hyperactivated in ΔG3BPs cells and known for their pro-apoptotic activity (Gannon et al., 2018; Li et al., 2017). Knocking out PKR or RNase L partially relieved dsRNA-triggered cell death in U2OSΔG3BPs cells, albeit not to the same extent as MAVS knock-out (Figures 5G & 5H). In A549ΔG3BPs cells, knocking out RNase L significantly rescued cell viability, but knocking out PKR did not (Figures S5D & S5E), suggesting that the contribution of each dsRNA-sensing pathway can vary depending on cell type. Another note-worthy observation was that, even in the same cell type, the effect of PKR knock-out was highly context-dependent; in the wild-type U2OS background, knocking out PKR increased dsRNA-triggered cell death (likely due to the inhibition of SG formation) (Figure 4E), while in the G3BPs-deficient background, knocking out PKR decreased dsRNA-triggered cell death (likely due to the relief of translational inhibition) (Figure 5G). This highlights the context-dependent impact of PKR on the life-death decision.

Altogether, our data indicate that SGs enable cells to avoid unnecessary dsRNA-mediated cell death by suppressing innate immune responses mediated by the RLR-MAVS-TNFα, PKR-eIF2α and OAS-RNase L pathways. The IRF3-mediated type I IFN response does not play a role, likely due to the strong negative feedback regulation through apoptotic caspases (Figure 5I).

### SGs suppress RLR signaling during viral infection, while restricting viral replication independent of RLRs

We next investigated how SGs impact innate immune responses and cell viability during viral infection. We used four viruses: Sendai virus (SeV), influenza A virus (IAV), vesicular stomatitis virus (VSV) and encephalitis myocarditis virus (EMCV). For IAV and VSV, the NS1-deletion variant of IAV (IAV^ΔNS1^) and the M51R variant of VSV (VSV^M51R^) were used as they were known to be more potent activators of innate immune pathways (Ahmed et al., 2003; Garcia-Sastre et al., 1998). SeV, IAV^ΔNS1^ and VSV^M51R^ predominantly activate RIG-I, whereas EMCV activates MDA5 (Loo et al., 2008).

SeV infection robustly induced SG formation and RLR signaling in WT U2OS cells (Figures 6A-C), but ΔG3BPs cells mounted significantly more efficient RLR signaling, as measured by the level of secreted IFNβ, IL-6, RANTES and TNFα or by *IFNβ* mRNA (Figures 6B and 6C). We also found that ΔG3BPs cells undergo more pronounced cell death than WT cells upon infection (Figure 6D), and that cell death in ΔG3BPs was largely dependent on MAVS, TNFα and caspases (Figures 6D and 6E), as was the case with *in vitro* transcribed dsRNA.

Analysis of EMCV, IAV^ΔNS1^ and VSV^M51R^ also showed similar results. That is, ΔG3BPs cells (whether U2OS or A549) showed heightened RLR signaling and more pronounced cell death than did WT cells in all conditions tested (Figures 6F and 6G; see also Figures S6A-G for more comprehensive analysis). Once again, RLR signaling and cell death were predominantly dependent on MAVS. These results suggest that the functions of SGs in suppressing RLR signaling and immune toxicity are preserved during infection with a broad range of viruses.

To examine how SGs impact viral replication and propagation, we measured cell-to-cell spreading and viral protein level per infected cell using immunofluorescence with antibodies against SeV, IAV^ΔNS1^ and VSV^M51R^ proteins (Figure 6H, antibodies against EMCV proteins were not available). With all three viruses, cell-to-cell spreading was more restricted in ΔG3BPs than in WT cells, and that this effect of G3BPs required MAVS (Figures 6I and S7A for SeV; S7B for VSV^M51R^; S7C for IAV^ΔNS1^). This result is consistent with the notion that hyper-active RLR signaling in ΔG3BPs cells restricts viral spreading. Intriguingly, the level of viral proteins per infected cell was higher in ΔG3BPs than in WT cells for all three viruses (Figures 6J and S7A for SeV; S7B for VSV^M51R^; S7C for IAV^ΔNS1^), and this effect of G3BPs was independent of MAVS. In keeping with this, we also observed a slight increase in overall viral mRNAs in ΔG3BPs cells and an additional increase in ΔG3BPs/ΔMAVS cells in most cases (Figures S7D-G). These data thus suggest that SGs have at least two independent functions during infection: (1) suppressing RLR signaling and consequent cell death, and (2) restricting viral replication independent of RLR signaling (Figure 6K, see Discussion).

### SGs protect cells from self-derived dsRNA accumulated under the ADAR1 deficiency

Our findings above showed that SGs have immune-suppressive and cell-protective roles against exogenous dsRNA regardless of the specific origins of dsRNA. We next asked whether these functions of SGs can be extended to cellular responses to self-derived dsRNA, which can erroneously accumulate under pathologic conditions. One such condition is the deficiency of RNA-editing enzyme ADAR1, which converts adenosine within dsRNA to inosine and disrupts duplex RNA structure (Walkley and Li, 2017). Previous studies showed that ADAR1 deficiency leads to accumulation of endogenous dsRNAs, aberrant activation of MDA5, PKR and OASes and ultimately pathogenesis of autoinflammatory diseases (Ahmad et al., 2018; Chung et al., 2018; Liddicoat et al., 2015; Mannion et al., 2014; Pestal et al., 2015; Rice et al., 2010; Wang et al., 2004). It was also shown that the immune effect of the ADAR1 deficiency is more evident upon IFNβ treatment (priming), which upregulates the levels of all three types of dsRNA sensors and further sensitizes cells to self-derived dsRNAs (Ahmad et al., 2018; Chung et al., 2018; Wang et al., 2004). Consistent with these reports, priming with IFNβ was necessary for robust SG formation and RLR-MAVS signaling (as measured by *IFNβ* mRNA induction) upon ADAR1 knock-down in WT cells (Figures 7A & 7B). Note that IFNβ priming without ADAR1 knock-down does not activate RLR signaling, in line with the ligand requirement for RLR signaling. In ΔG3BPs cells, ADAR1 knock-down resulted in more potent RLR signaling than in WT cells, both in the presence and absence of IFNβ pre-treatment (Figure 7B). ADAR1 knock-down also led to significantly increased cell death in ΔG3BPs than in WT cells (Figure 7C). As with exogenous dsRNA stimulation, this cell death was predominantly dependent on MAVS and partly on PKR and RNase L (Figure 7C), and could be relieved by anti-TNFα and pan-caspase inhibitor (Figure 7D). As with ΔG3BPs cells, ΔUBAP2L cells were also hypersensitive to ADAR1 knock-down than WT cells (Figure 7E), further supporting the notion that SG deficiency increases sensitivity to ADAR1 deficiency (Figure 7E). These results together suggest that SGs suppress excessive immune activation and cell death, whether they are triggered by exogenous or endogenous dsRNA.

**Figure 7.**
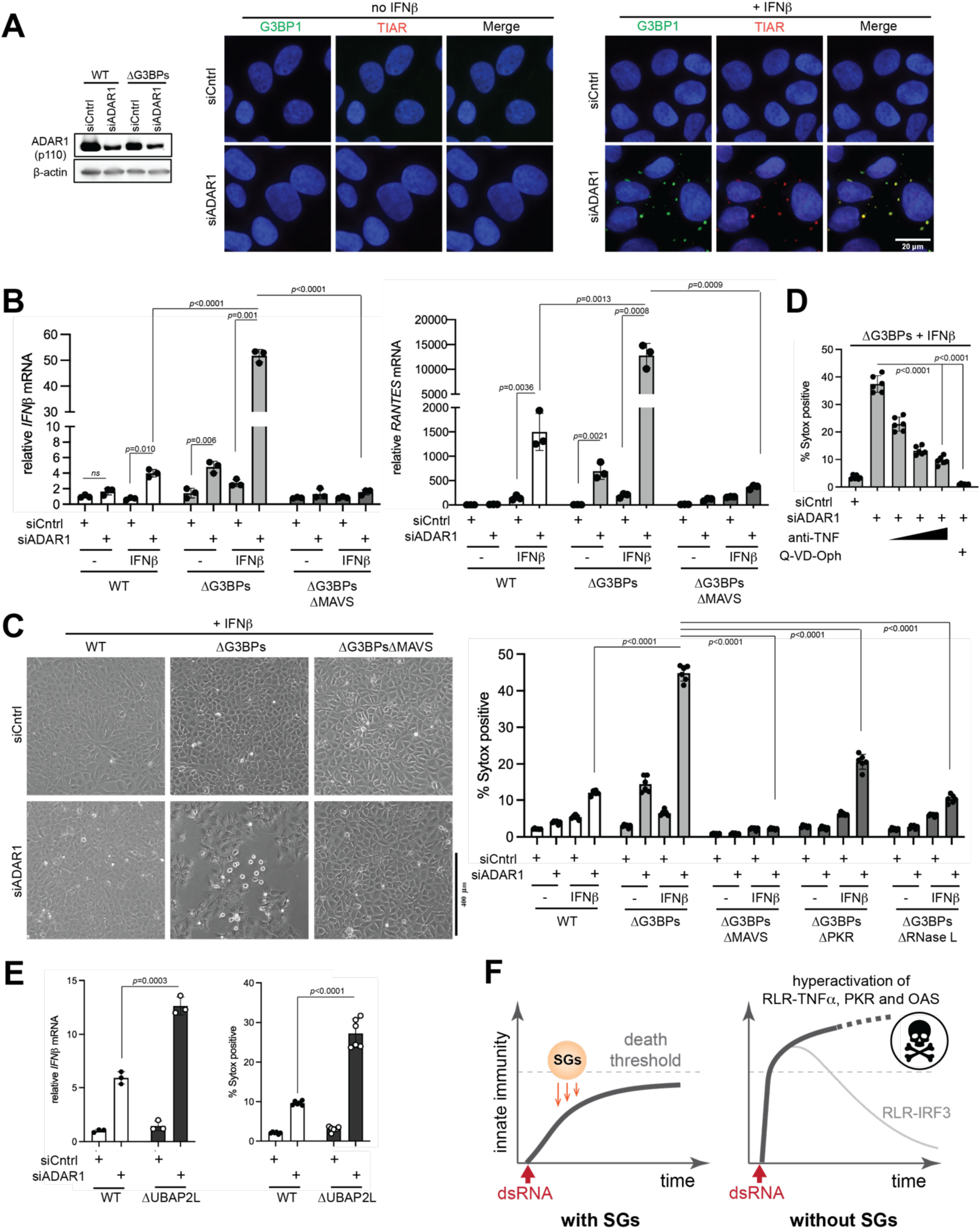
SGs suppress immune response to self-derived dsRNAs under the ADAR1 deficiency. **A.** Immunofluorescence (IF) analysis of G3BP1 (green) and TIAR (red) in WT U2OS cells in the presence or absence of ADAR1 knock-down and IFNβ priming. Cells were transfected with siADAR1 (or siCntrl) for 24 hrs and then treated with IFNβ (10 ng/ml) for additional 24 hrs prior to imaging. **B.** Antiviral signaling upon ADAR1 knock-down, as measured by the level of *IFNβ* mRNAs. WT, ΔG3BPs and ΔG3BPs/ΔMAVS U2OS cells were treated with IFNβ and siADAR as in (A). **C.** Cell death upon ADAR1 knock-down, as measured by brightfield images (left) and Sytox uptake (right). U2OS cells were treated with IFNβ and siADAR as in (A). **D.** Effect of anti-TNF and pan-caspase inhibitor (Q-VD-Oph) on cell death upon ADAR1 knock-down, as measured by Sytox uptake. U2OSΔG3BPs cells were treated with IFNβ and siADAR as in (A). **E.** Antiviral signaling and cell death upon ADAR1 knock-down in WT vs. ΔUBAP2L U2OS cells. Experiments were performed as in (A-C). All samples were treated with IFNβ (10 ng/ml). **F.** Schematic summarizing the roles of SGs in protecting cells from dsRNA. SGs suppress a broad range of dsRNA-triggered innate immune pathways (RLR, PKR and OASes), regardless of the origin of dsRNA. This may enable cells to increase the temporal window for mounting an appropriate immune response while maintaining its magnitude below the “death” threshold. In the absence of SGs, dsRNA sensing pathways are hyperactivated, leading to an excessive innate immune response and consequent cell death. The IRF3-IFN axis downstream of RLR-MAVS does not contribute to cell death and often displays a dynamic temporal behavior characterized by a sharp peak followed by a strong decline due to caspase-dependent feedback regulation. Data are presented in means ± SD. *p* values were calculated using two-tailed unpaired Student’s t test (ns, *p*>0.05). All data are representative of three independent experiments.

## Discussion

Phase separation has recently emerged as a widespread phenomenon that occurs in many biological processes, from transcription to signal transduction (Alberti and Hyman, 2021; Hnisz et al., 2017). Functions of phase separation, however, remain unclear in most cases. SGs are one such phase separated entities with poorly characterized functions. We here use two independent SG-deficient genetic backgrounds that lack key SG nucleators, G3BP1/2 and UBAP2L, to show that SGs exert a negative impact on dsRNA-dependent innate immune pathways. RLRs, PKR and OASes and downstream effector molecules were highly enriched within SGs, but dsRNA itself was not, arguing against the idea that SGs are sites of innate immune activation or that SGs selectively recruit activated molecules. Rather, these innate immune molecules appear to be recruited to SGs independent of their activation state and their sequestration within SGs may prevent their activation. We found that absence of SGs leads to overactivation of the RLR pathway, excessive production of TNFα and other pro-apoptotic genes, and consequent cell death. The SG deficiency also results in hyperactivation of PKR and OASes, further contributing to dsRNA-dependent cell death. We thus propose that SGs function as a “buffer” or “shock absorber” to enable the controlled activation of innate immune pathways upon dsRNA introduction, by increasing the temporal window for mounting an appropriate immune response while maintaining its magnitude below the “death” threshold (Figure 7F). Given our findings that SGs control cellular response not only to viral dsRNA, but also to self-derived dsRNAs that can be generated from many dysregulated cellular processes, SGs may serve as a key immune modulator in a broad range of pathophysiological conditions.

How can we reconcile our findings with previous reports suggesting that SGs amplify RLR signaling? One potential explanation may be in the complexity with which cells regulate IRF3-dependent IFN induction, one branch of the RLR pathways that is often used as a measure of the overall RLR signaling activity. We found that unlike other RLR-dependent signaling axes (*e.g.* those inducing TNFα or RANTES), the IRF3-IFNs axis in SG-deficient cells often responds to dsRNA with an initial spike followed by a marked decline. This decline, which was seen only in SG-deficient cells, is a result of strong negative feedback regulation by apoptotic caspases that are activated in SG-deficient cells. We speculate that such feedback regulatory mechanism for the IRF3-IFNs axis may account for seemingly conflicting data in the literature. Additionally, specific methods of SG inhibition may have further contributed to the confusion. Unlike depleting SG nucleators (*e.g.* G3BPs and UBAP2L), we found that depletion of PKR, which also impairs dsRNA-triggered SG formation, results in a more complex, time-dependent signaling behavior. While RLR signaling in ΔPKR cells at early time point was similar to that of ΔG3BPs and ΔUBAP2L cells, at later time points ΔPKR cells showed a divergent behavior. This likely reflects the fact that PKR regulates both translation and SG formation, while G3BPs and UBAP2L only affect SGs without altering translational control. Additionally, PKR is not only an upstream inducer of SGs, but is also subject to SGs-mediated feedback regulation. These complex relationships between PKR and SGs caution using ΔPKR as the sole genetic model for the SG-deficiency.

Our results also show a dual role of SGs in antiviral immunity – suppressing RLR-mediated excessive inflammation and restricting viral replication independent of the RLR pathway. Previous studies showed that many viral RNAs and proteins are localized within SGs, which may exert sequestration and inhibitory functions independent of RLRs (Emara and Brinton, 2007; Jayabalan et al., 2021). Additionally, all viruses rely on the host ribosome to synthesize viral proteins, but the 40S subunit is sequestered within SGs (Kedersha et al., 2002; Kimball et al., 2003), which could further limit the translational resource for viruses. Our observations thus highlight the multi-functional nature of SGs that cannot be simply categorized into anti- or pro- viral activities. This is in line with the observation that different viruses cope with SGs differently; some inhibit SGs, while others alter or take advantage of SGs (Feng et al., 2014; McCormick and Khaperskyy, 2017; White and Lloyd, 2012). The diverse functions of SGs may instead be understood as a cellular mechanism for maintaining cell homeostasis – a common consequence of dampening the toxic immune response and viral replication. Thus, our study provides a new foundation to investigate the role of phase separation in host-virus interactions, dsRNA biology and innate immune regulation, with the potential of guiding new avenues for antiviral or immune-modulatory therapies.

## Acknowledgement

We acknowledge Drs. Paul Anderson, Igor Brodsky and the rest of the Hur lab members for helpful discussions. We thank Dr. Takashi Fujita for sharing anti-IRF3 antibody. Cell images for z-stack analysis were collected at the Nikon imaging center at Harvard Medical School. This work was supported by NSF fellowship (CC), Roche post-doctoral fellowship (HW), and NIH grants to SML (R00GM124458), PI (R01GM126150), BT (R01 AI145882, R21 AI171083, R01 AI170487), and SH (R01AI154653, R01AI111784, DP1AI152074). SH is a Howard Hughes Medical Institute investigator.

## Data Availability

Raw data from RNA-seq were deposited in the Gene Expression Omnibus (http://www-ncbi-nlm-nih-gov.ezp-prod1.hul.harvard.edu/geo/) under accession number GSE173953.

## Code availability

Publicly available codes were used for RNA-seq analysis. See reporting summary.

## Biological material availability

All unique materials will be available upon publication and request.

## STAR*Methods

### Key resources table

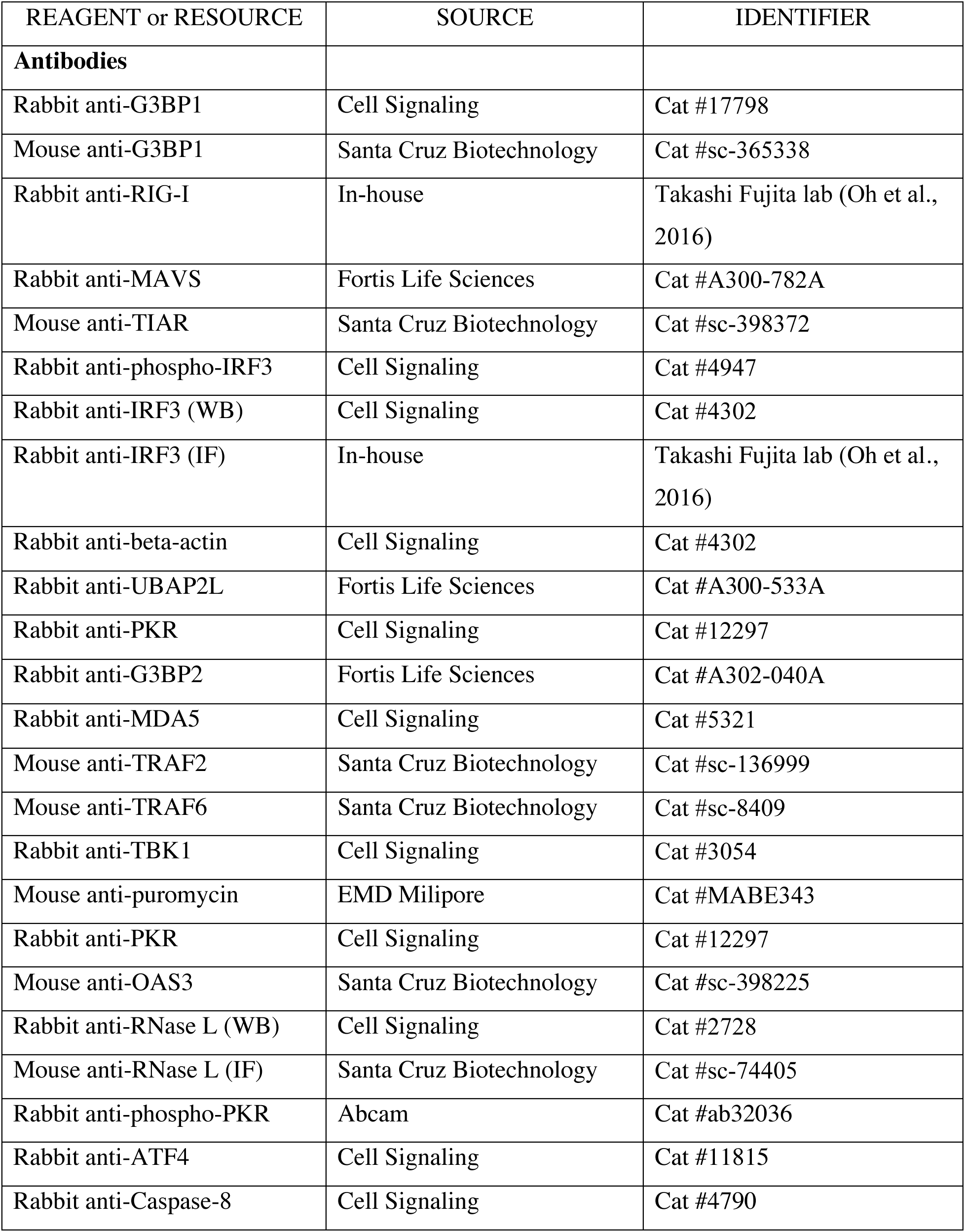
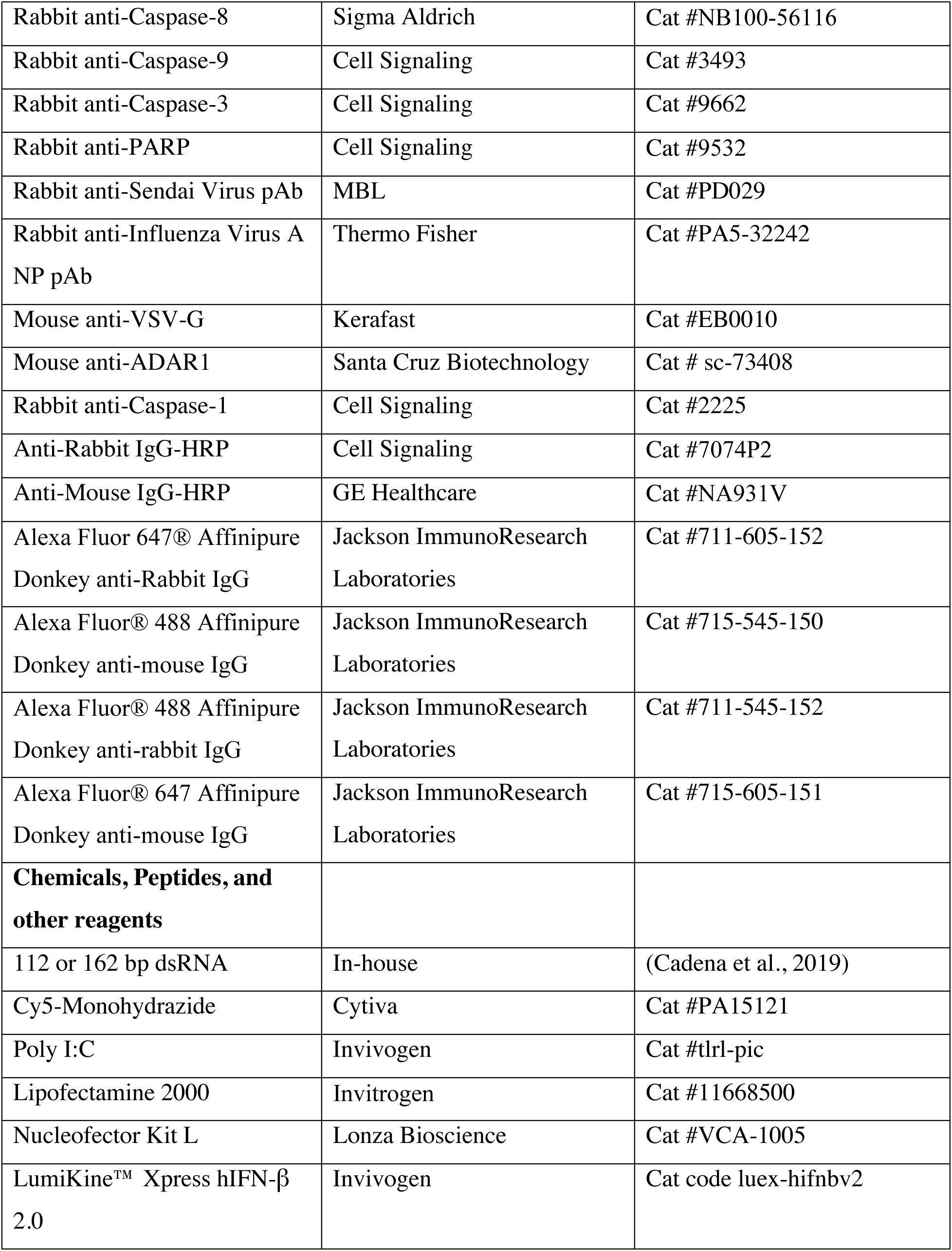
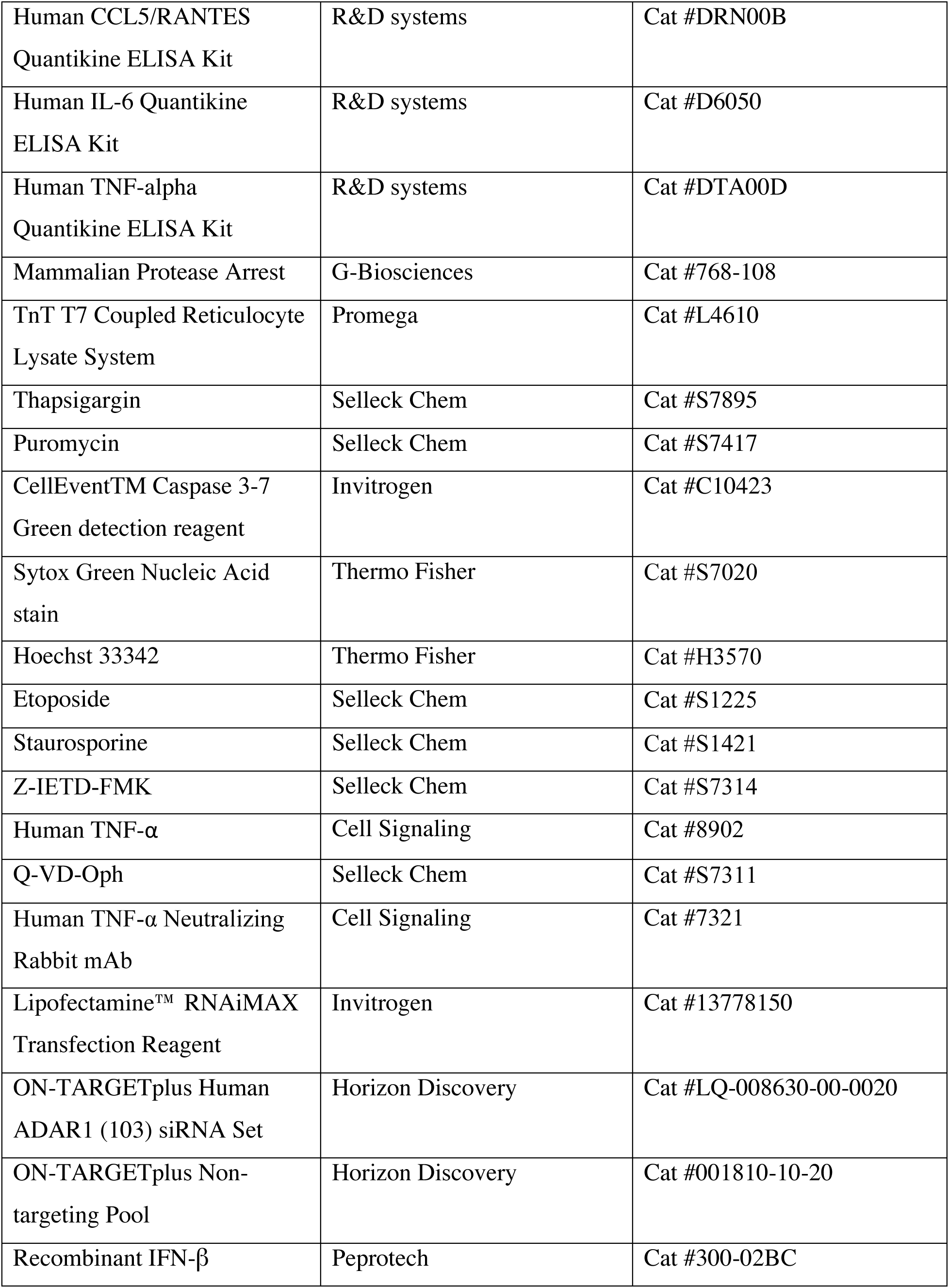
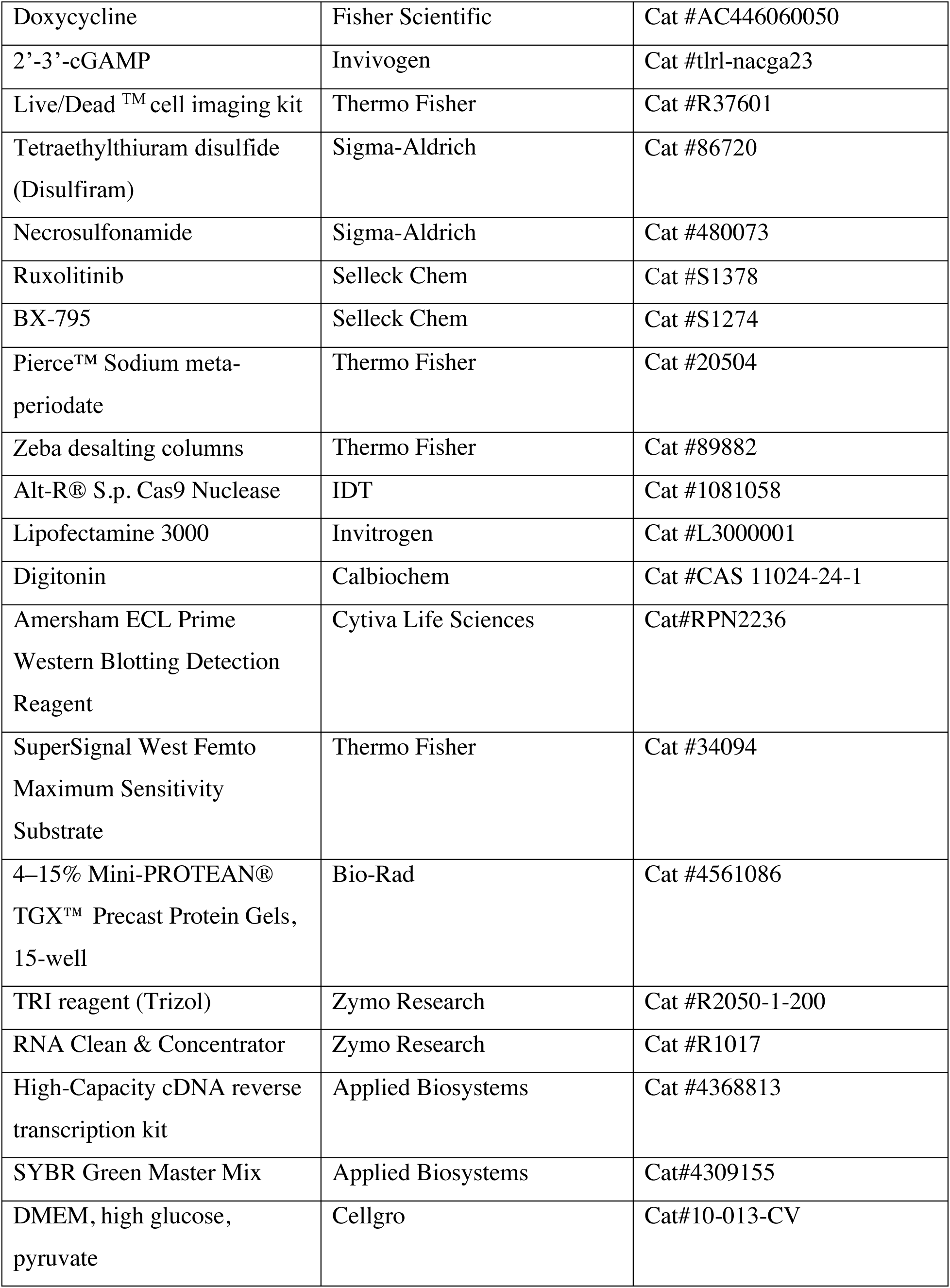
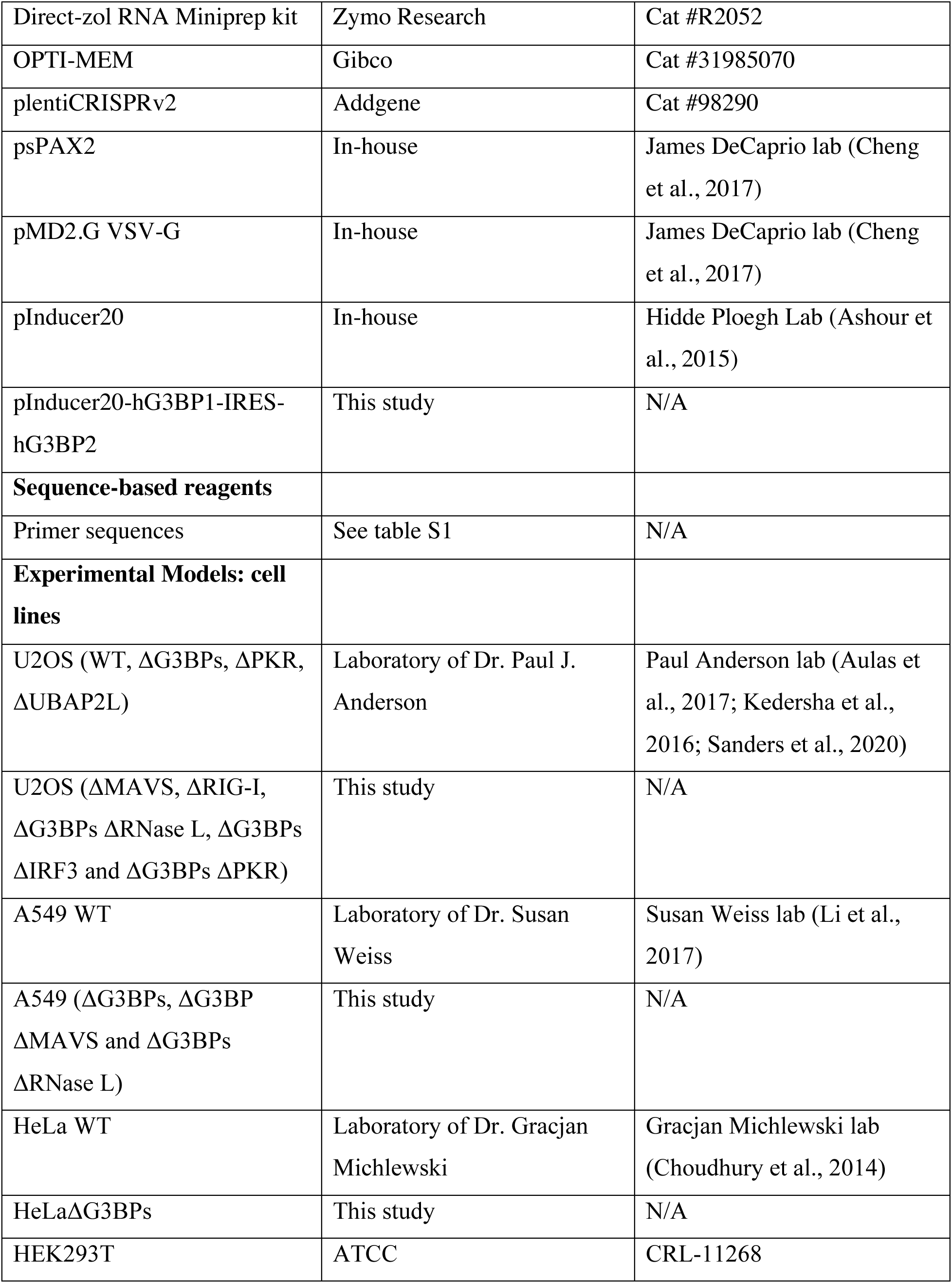
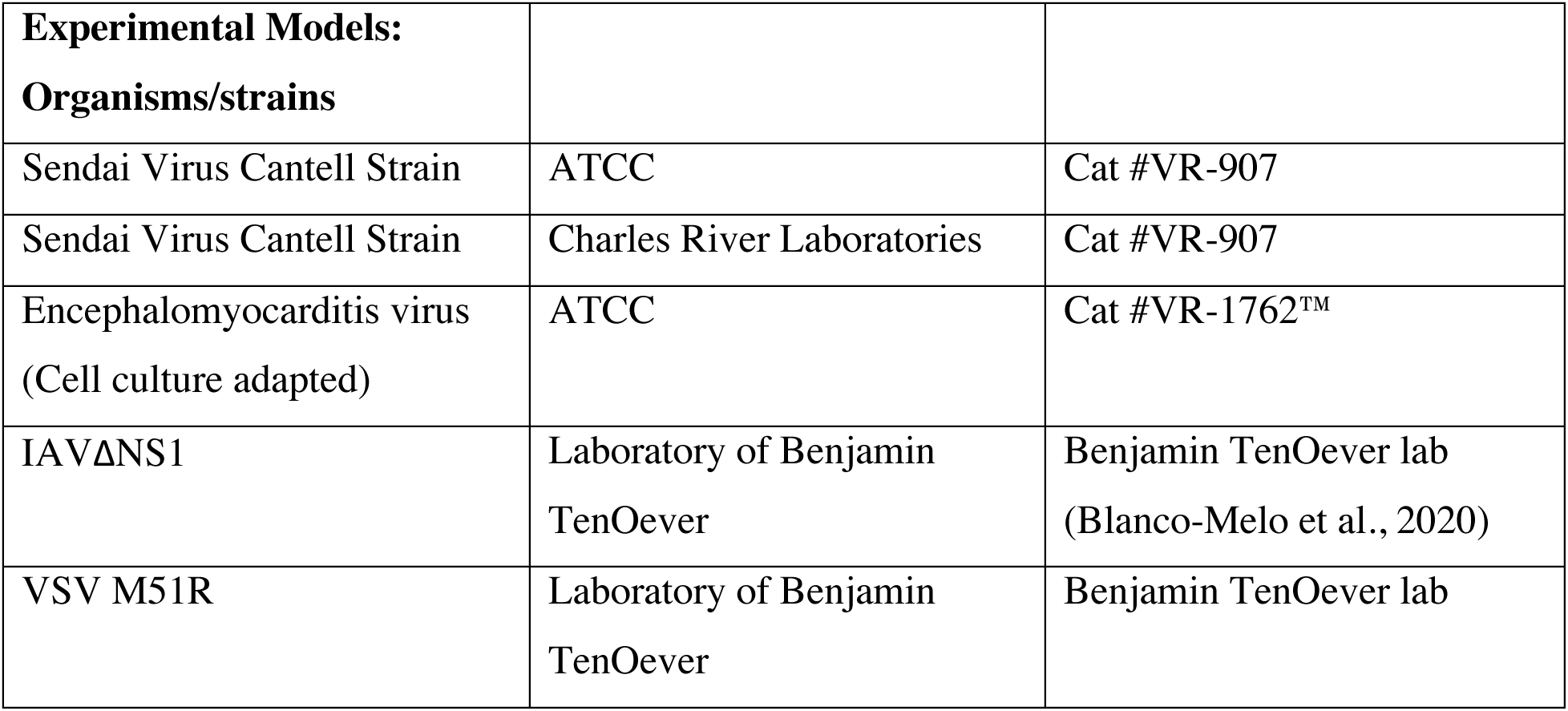

### Contact for reagent and resource sharing

Further information and requests for reagents may be directed to and will be fulfilled by the corresponding author Sun Hurs.

### Experimental Model and Subject Details Cell Lines

#### U2OS cells

Cells were maintained in DMEM (High glucose, L-glutamine, Pyruvate) with 10% fetal bovine serum

#### A549 cells

Cells were maintained in DMEM (High glucose, L-glutamine, Pyruvate) with 10% fetal bovine serum

#### HEK293T cells

Cells were maintained in DMEM (High glucose, L-glutamine, Pyruvate) with 10% fetal bovine serum

#### HeLa cells

Cells were maintained in DMEM (High glucose, L-glutamine, Pyruvate) with 10% fetal bovine serum

---

**Figure S1.**
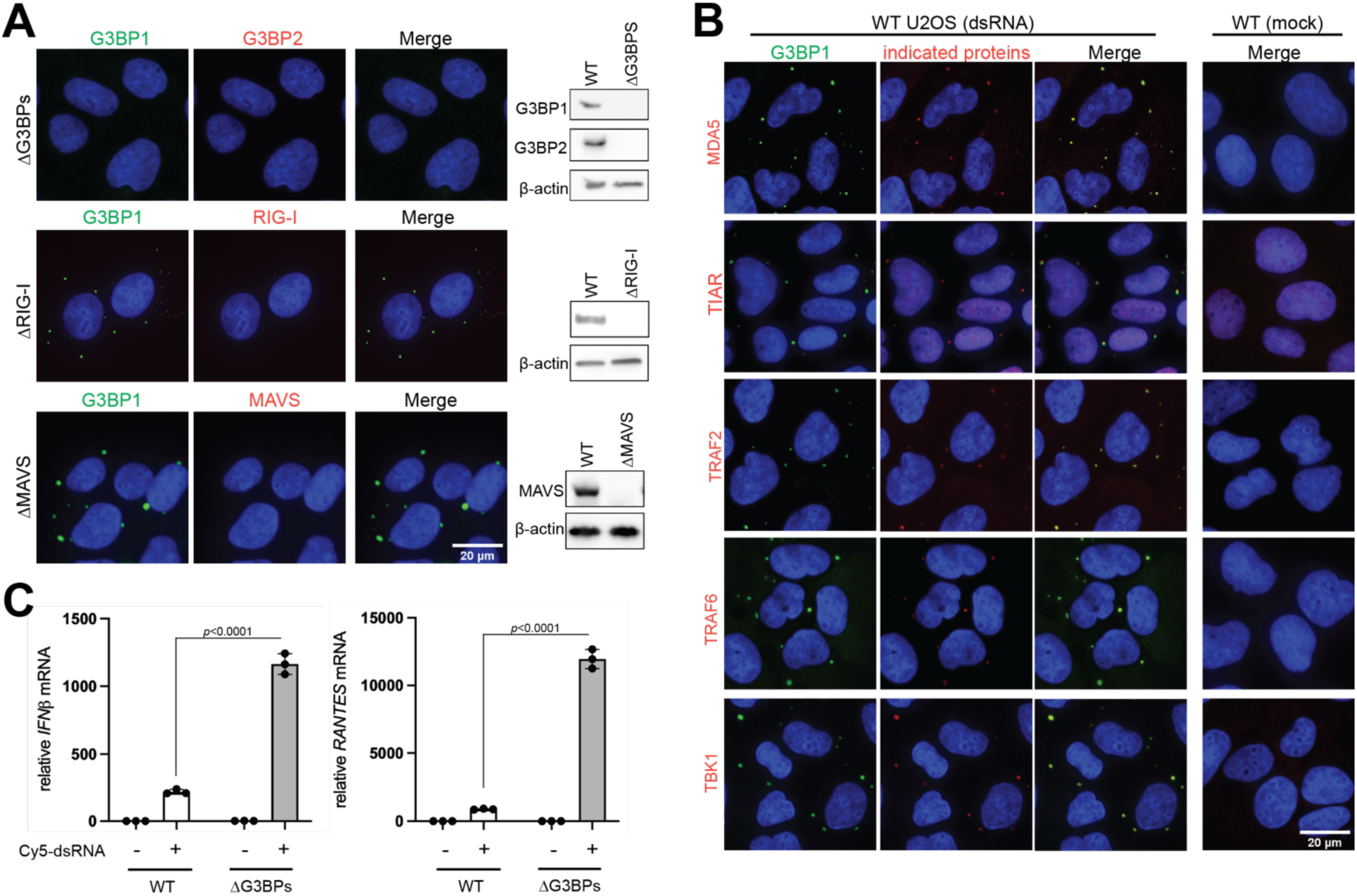
dsRNA-triggered SGs are enriched for innate immune signaling molecules in the RLR pathway. Related to Figure 1. **A.** Validation of antibodies for G3BP1, RIG-I and MAVS using individual knockout U2OS cells. Cells were transfected with 162 bp dsRNA (500 ng/ml) and imaged at 6 hr post-dsRNA as in Figure 1B. **B.** Immunofluorescence analysis of MDA5, TIAR, TRAF2, TRAF6 and TBK1 (red) with G3BP1 (green) at 6 hr post-dsRNA in U2OS cells. **C.** Antiviral signaling in U2OS cells (WT vs ΔG3BPs), as measured by the level of *IFNβ* (left) and *RANTES* (right) mRNAs. Cells were transfected with 162 bp dsRNA 3’-labeled with Cy5 (500 ng/ml) for 6 hrs. 162 bp dsRNA activates the RLR pathway more efficiently in ΔG3BPs cells than in WT cells, regardless of the presence of 3’-Cy5 label.

**Figure S2.**
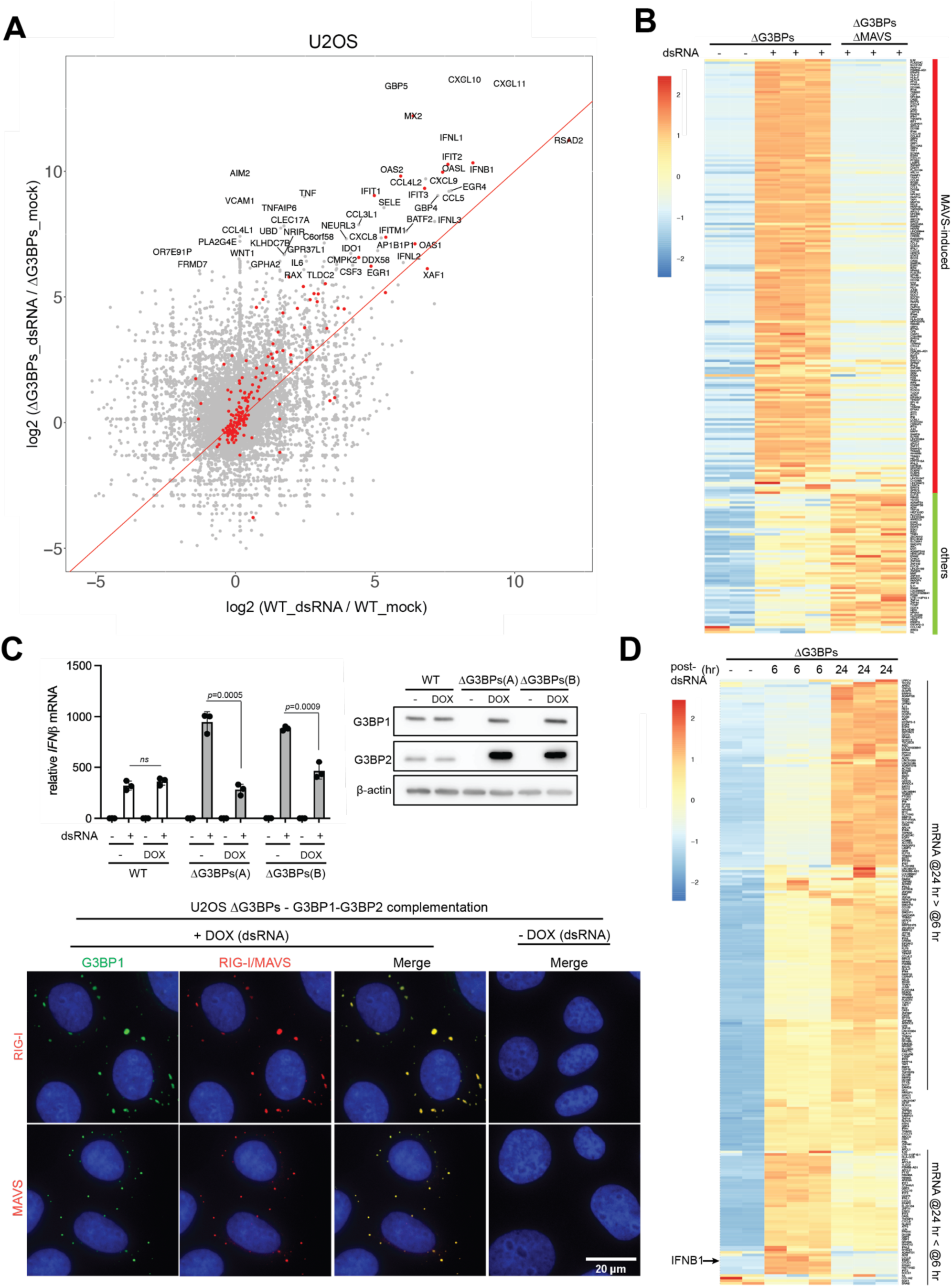
RLR signaling is hyperactive in SG-deficient ΔG3BPs U2OS cells. Related to Figure 1. **A.** Scatter plot of log2-fold change (lfc2) upon dsRNA stimulation for 6 hrs. Lfc2 values in U2OS WT cells were plotted against those in ΔG3BPs cells. Differentially expressed genes with lfc2 > 6 in ΔG3BP cells are labeled. Genes annotated as “type I interferon production” (GO:0032606) and “response to type I interferon” (GO:0034340) are colored red. **B.** Heat map of z-scores comparing the levels of mRNA in U2OS cells (ΔG3BPs vs. ΔG3BPs/ΔMAVS cells with or without dsRNA stimulation for 6 hrs). Genes showing lfc2 >2 (with *p*_adj<0.05) upon dsRNA stimulation in WT cells were shown. **C.** G3BPs complementation assay. Relative levels of *IFNβ* mRNA in U2OSΔG3BPs cells with and without complementation of G3BP1/2, which were expressed under the control of a doxycycline (DOX)-inducible promoter. Two separate clones (A and B) were treated with DOX (1 μg/ml) for 24 hrs and transfected with 162 bp dsRNA (500 ng/ml) for 6 hrs prior to analysis. Bottom: IF analysis of SGs with and without DOX induction. **D.** Heat map of z-scores comparing the levels of mRNA in U2OSΔG3BPs cells at 6 and 24 hr post-dsRNA; indicated genes were from (B) and were re-ordered by hierarchical clustering. Data are presented in means ± SD. *p* values were calculated using two-tailed unpaired Student’s t test (ns, *p*>0.05). All data are representative of two to three independent experiments. Images were taken from representative field of view. Raw data for the heatmaps can be found in the supplemental files (data 2 and 3).

**Figure S3.**
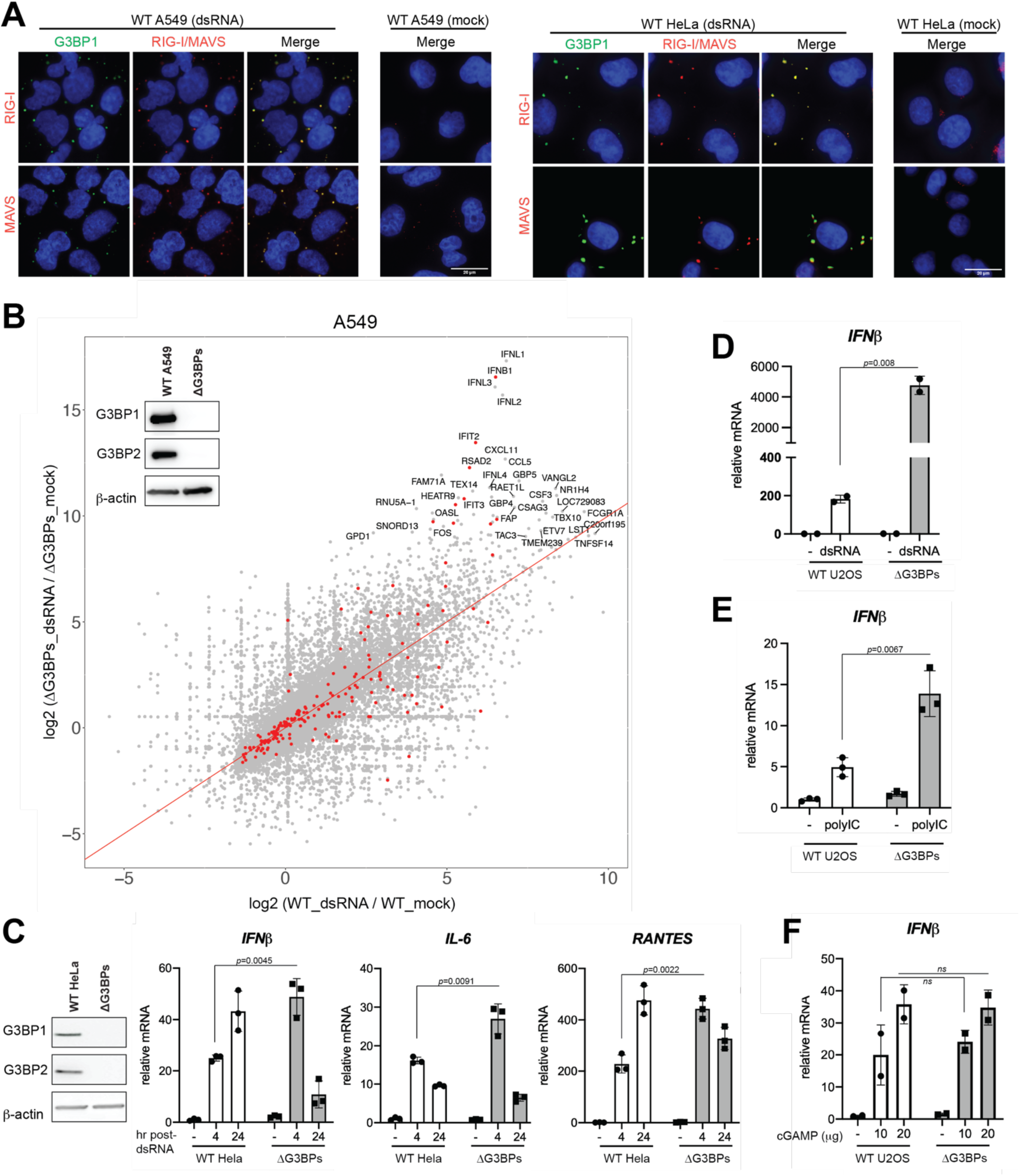
G3BPs suppress RLR signaling in both A549 and HeLa cells. Related to Figure 1. **A.** Immunofluorescence analysis of RIG-I (red), MAVS (red) and G3BP1 (green) in A549 cells (left) and HeLa cells (right). Cells were transfected with 162 bp dsRNA containing 5’ppp (500 ng/ml) for 6 hrs. Nuclei were stained with Hoechst 3342. **B.** Scatter plot of log2-fold change (lfc2) upon dsRNA stimulation (500 ng/mL) for 4 hrs. Lfc2 values in WT A549 were plotted against those in ΔG3BPs cells. Differentially expressed genes with lfc2 > 9 in ΔG3BP cells were labeled. Genes annotated as “type I interferon production” (GO:0032606) and “response to type I interferon” (GO:0034340) were colored red. **C.** Antiviral signaling in HeLa cells (WT vs ΔG3BPs), as measured by the level of *IFNβ* (left), *IL-6* (middle), and *RANTES* (right) mRNAs. Cells were transfected with 162 bp dsRNA with 5’ppp (500 ng/ml) for 4 or 24 hrs. **D.** Antiviral signaling in U2OS cells (WT vs ΔG3BPs) in response to dsRNA electroporation, as measured by the level of *IFNβ* mRNAs at 8 hr post-electroporation. **E.** Antiviral signaling in U2OS cells (WT vs ΔG3BPs) in response to poly I:C transfection, as measured by the level of *IFNβ* mRNAs at 6 hr transfection. **F.** Antiviral signaling in U2OS cells (WT vs ΔG3BPs) in response to 2′3′-Cyclic GMP-AMP (cGAMP, 10 or 20 μg), as measured by the level of *IFNβ* mRNAs at 6 hr post-treatment. Data are presented in means ± SD. *p* values were calculated using two-tailed unpaired Student’s t test (ns, *p*>0.05). RNA-seq data were confirmed by RT-qPCR analysis of select few genes in three independent experiments. All other data are representative of three independent experiments. Images were taken from representative field of view.

**Figure S4.**
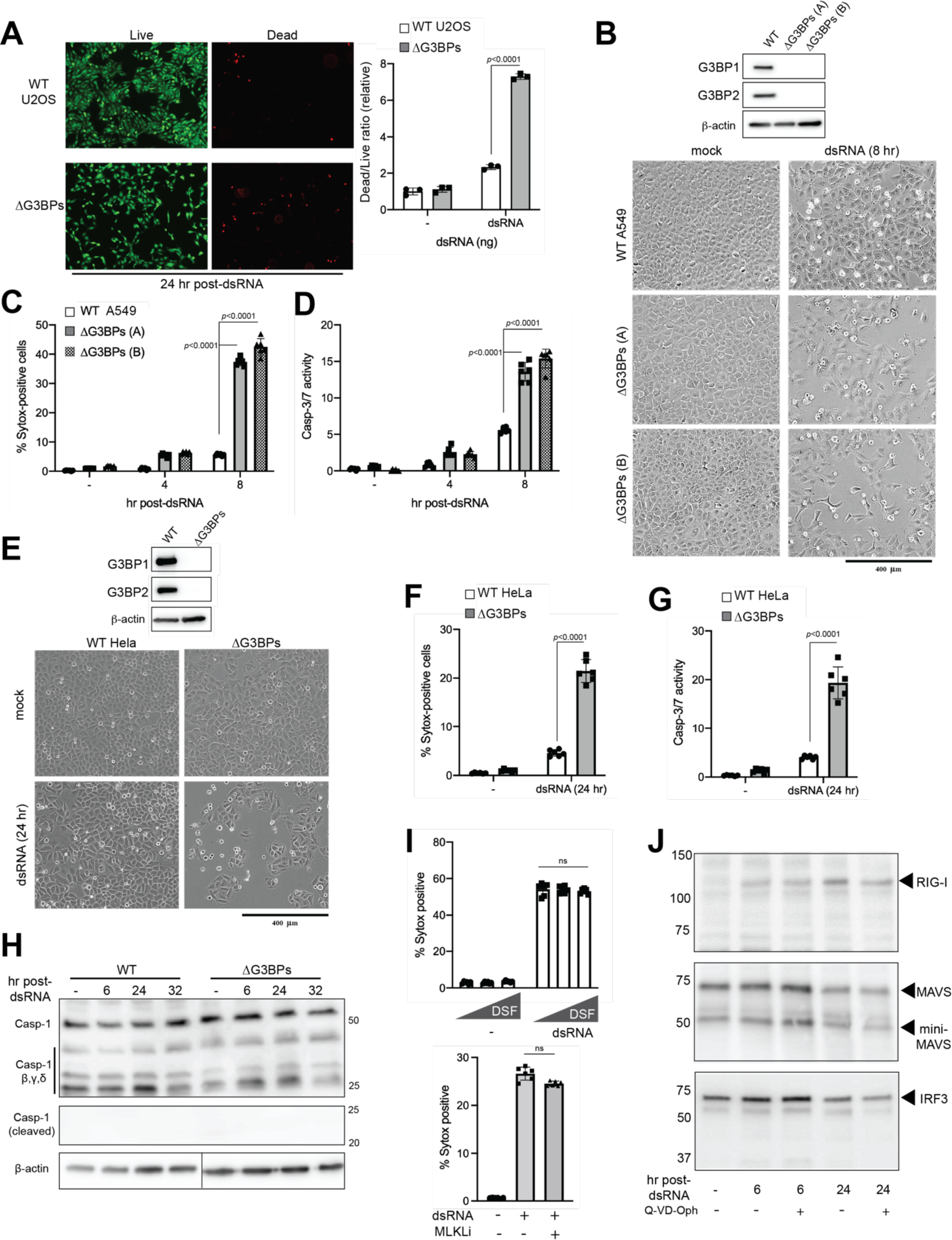
SG deficiency results in more pronounced apoptosis in U2OS, A549 and HeLa cells upon dsRNA stimulation. Related to Figure 4. **A.** Cell death in WT vs. ΔG3BPs U2OS cells in response to 162 bp dsRNA transfection (500 ng/ml). Cells were stained with the LIVE/DEAD® Viability/Cytotoxicity stain (Thermo) 24 hr post-dsRNA and were imaged using a Nikon Eclipse TS2R inverted microscope (left). The ratio of dead-to-live cells were determined by the ratio of the DEAD stain intensity (red) to the LIVE stain intensity (green) as quantitated by ImageJ (right). **B-D.** Cell death in WT vs. ΔG3BPs A549 cells in response to 162 bp dsRNA transfection (500 ng/ml). Two independent clones of ΔG3BPs were analyzed. Cell death was measured at 8 hr post-dsRNA by brightfield microscopy (B), Sytox uptake (C), and caspase 3/7 activity assay (D). **E-G.** Cell death in WT vs. ΔG3BPs HeLa cells in response to 162 bp dsRNA transfection (500 ng/ml). Cell death was measured at 24 hr post-dsRNA by brightfield microscopy (E), Sytox uptake (F), and caspase 3/7 activity assay (G). **H.** Caspase-1 cleavage analysis of WT and ΔG3BPs U2OS cells. Cells were transfected with 162 bp dsRNA (500 ng/ml) and were analyzed at indicated time. No caspase-1 cleavage was observed in either WT or ΔG3BPs cells upon dsRNA transfection. **I.** Effect of the gasdermin D inhibitor (disulfiram, DFS) and MLKL inhibitor (necrosulfonamide, MLKLi) on dsRNA-induced cell death in U2OSΔG3BPs. Cells were pre-treated with DMSO, disulfiram (10 or 25 μM) or MLKLi (1 μM) and transfected with 162 bp dsRNA (500 ng/ml). **J.** Analysis of RIG-I, MAVS or IRF3 cleavage upon dsRNA stimulation. U2OSΔG3BPs cells were transfected with 162 bp dsRNA (500 ng/ml) in the presence or absence of pan-caspase inhibitor (Q-VD-OPh, 10 μM). Cells were harvested at indicated time points and analyzed by SDS-PAGE. Data are presented in means ± SD. *p* values were calculated using two-tailed unpaired Student’s t test (ns, *p*>0.05). All data are representative of two to three independent experiments.

**Figure S5.**
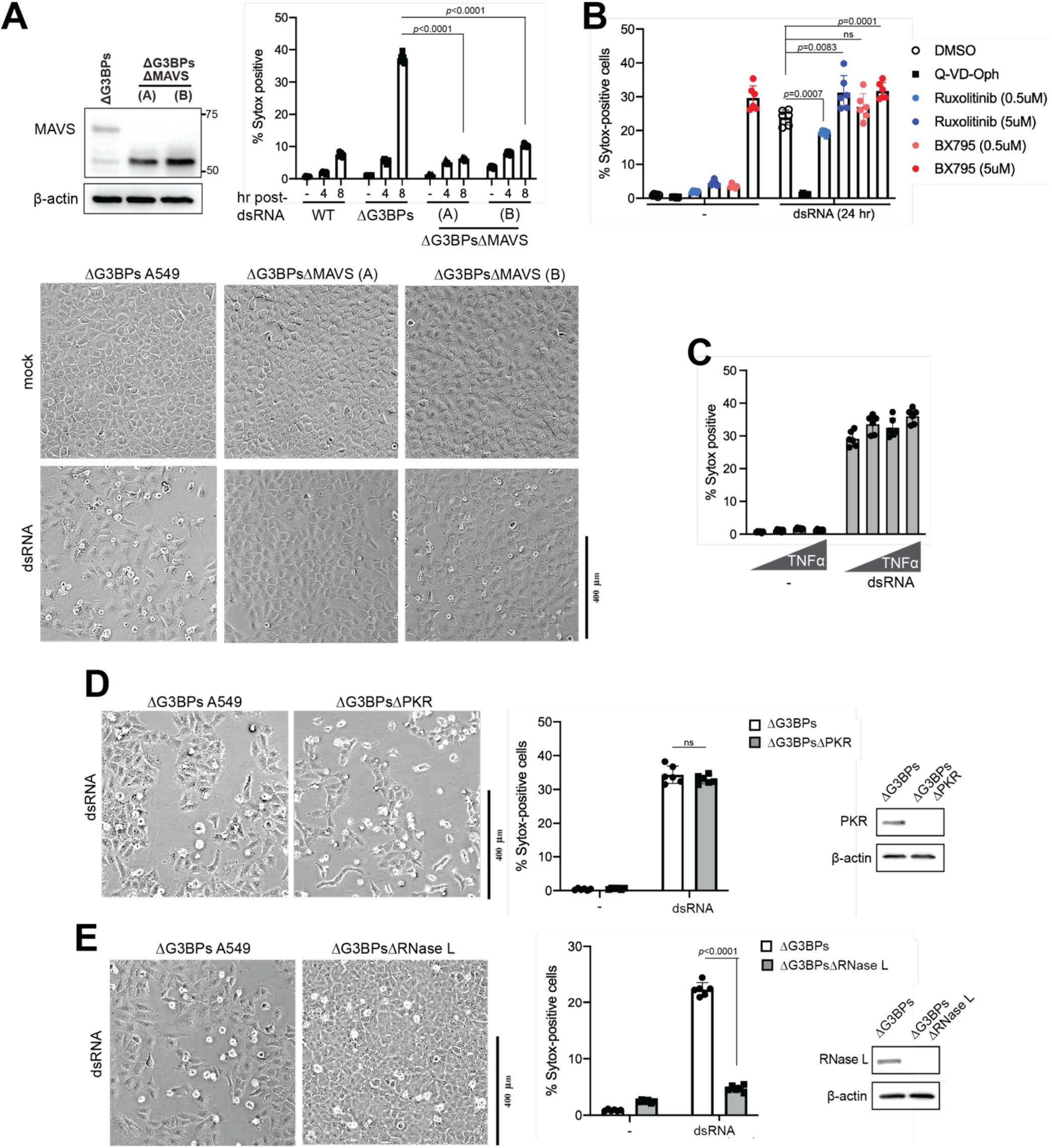
Apoptosis in SG-deficient ΔG3BPs cells is dependent on MAVS. Related to Figure 5. **A.** Cell death in ΔG3BPs and ΔG3BPsΔMAVS A549 as measured by Sytox uptake (top) and bright field microscopy (bottom) at 8 hr post-dsRNA. Cells were transfected with 162 bp dsRNA as in Figure 1B. **B.** Cell death in U2OSΔG3BPs cells in the presence of an inhibitor for TBK1 (BX795) or JAK (Ruxolitinib). Pan-caspase inhibitor Q-VD-Oph (10 μM) was used for comparison. Cells were pre-treated with the indicated inhibitor 30 min prior to dsRNA transfection (500 ng/ml). Cell death was measured by Sytox uptake at 24 hr post-dsRNA. **C.** Cell death in U2OSΔG3BPs cells in response to an increasing concentration of TNFα (10, 20, 50 ng/ml). Cell death was measured by Sytox uptake at 24 hr post-treatment. **D.** Cell death in ΔG3BPs and ΔG3BPsΔPKR A549 cells in response to dsRNA. Cell death was measured by brightfield microscopy (left) and Sytox uptake (right) at 8 hr post-dsRNA. **E.** Cell death in ΔG3BPs and ΔG3BPsΔRNase L A549 cells in response to dsRNA. Cell death was measured by brightfield (left) and Sytox uptake (rigjt) at 8 hr post-dsRNA. Data are presented in means ± SD. *p* values were calculated using two-tailed unpaired Student’s t test (ns, *p*>0.05). All data are representative of three independent experiments.

**Figure S6.**
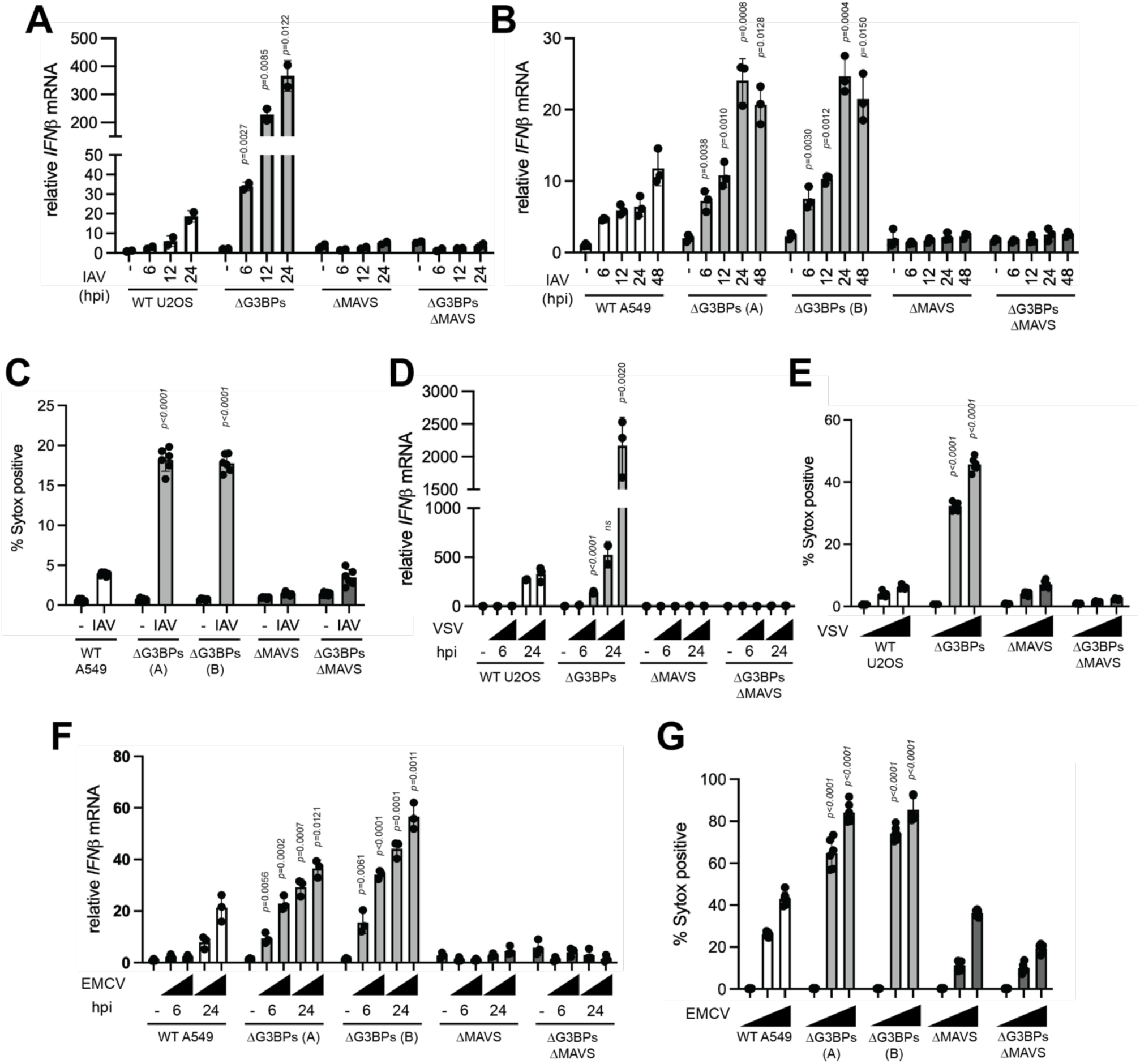
SGs suppress immune response and cell death during viral infection. Related to Figure 6. **A.** Antiviral signaling upon infection with IAV^ΔNS1^ (MOI=1) in U2OS cells, as measured by *IFNβ* mRNA. **B.** Antiviral signaling upon infection with IAV^ΔNS1^ (MOI=0.1) in A549 cells, as measured by *IFNβ* mRNA. **C.** Cell death upon infection with IAV^ΔNS1^ (MOI=0.1), as measured by Sytox uptake. A549 cells were infected virus as in (B) and analyzed at 24 hpi. **D.** Antiviral signaling upon infection with VSV^M51R^ (MOI=0.1 or 1) in U2OS cells, as measured by *IFNβ* mRNA. **E.** Cell death upon infection with VSV^M51R^ (MOI=0.1 or 1), as measured by Sytox uptake. U2OS cells were infected with VSV^M51R^ as in (D) and analyzed at 24 hpi. **F.** Antiviral signaling upon infection with EMCV (MOI=0.01 or 0.1) in A549 cells, as measured by *IFNβ* mRNA. **G.** Cell death upon infection with EMCV (MOI=0.1 or 0.3), as measured by Sytox uptake. A549 cells were infected with EMCV as in (F) and analyzed at 24 hpi. Data are presented in means ± SD. All data are representative of at least three independent experiments. *p* values were calculated using two-tailed unpaired Student’s t test (ns, *p*>0.05). Unless indicated otherwise, *p* values were calculated in comparison to WT values at equivalent conditions.

**Figure S7.**
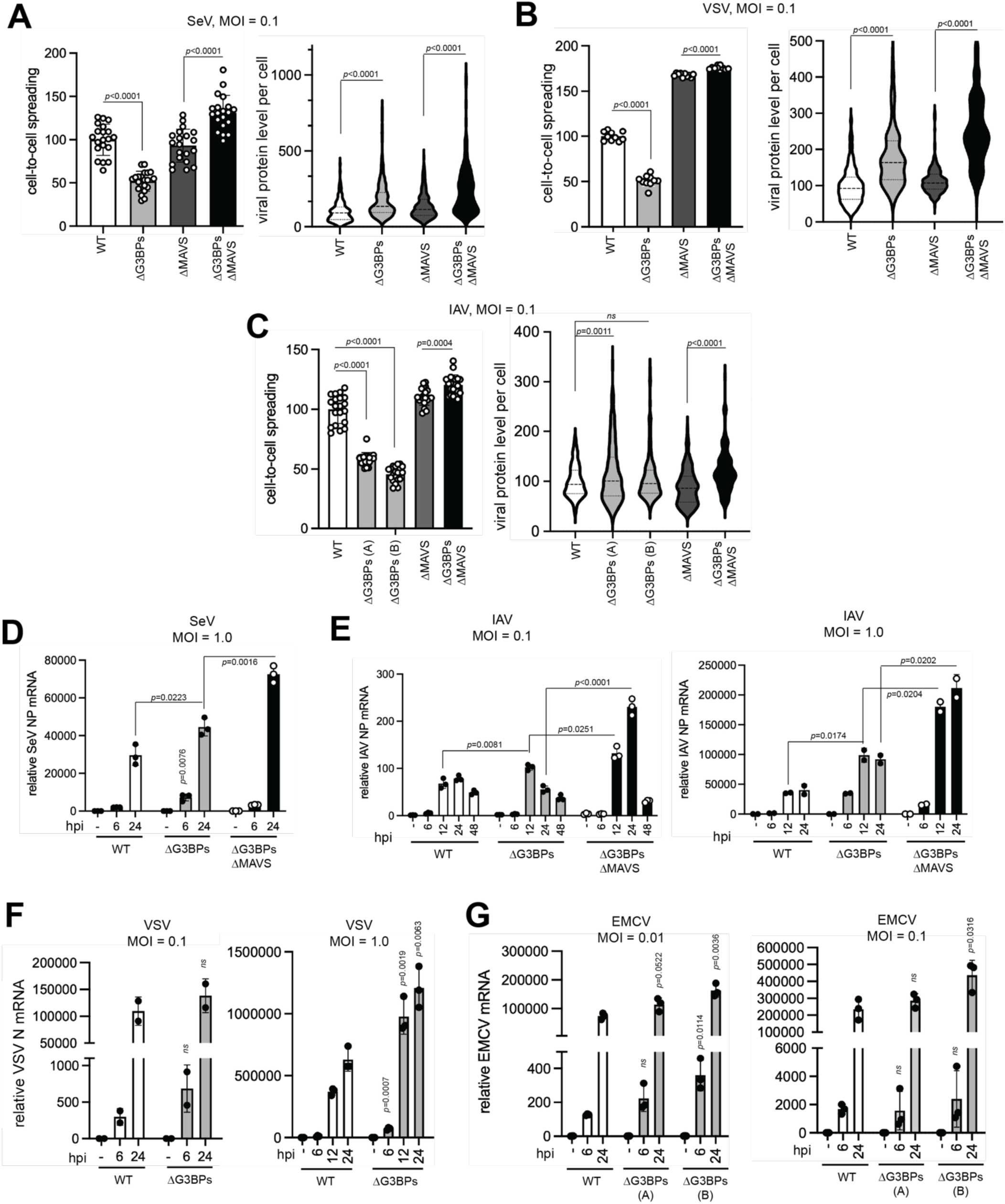
SGs also suppress viral replication independent of the RLR pathway. Related to Figure 6. **A.** SeV cell-to-cell spreading and viral protein level per infected cell. Left: Relative cell-to-cell spreading of SeV. U2OS cells were infected with SeV (MOI=0.1), stained with anti-SeV serum 18 hpi, analyzed for number of cells above the background fluorescence (i.e. mock infection staining) per field of view. Each data point represents a field of view (n=20). Right: Relative level of SeV protein staining in infected cells, as measured by corrected total cell fluorescence (CTCF). Each data point represents an infected cell (n=200). Data were normalized against the WT average value in both graphs. **B.** Same as (A) for VSV^M51R^ (MOI=0.1) infection in U2OS cells. (Left) n=10, (Right) n=200. **C.** Same as (A) for IAV^ΔNS1^ (MOI=0.1) infection in A549 cells. (Left) n=20, (Right) n=200. **D.** Level of SeV NP mRNA during infection. U2OS cells were infected with SeV (MOI=1) and analyzed at indicated time points. **E.** Level of IAV^ΔNS1^ NP mRNA during infection. Left: A549 cells were infected with IAV^ΔNS1^ (MOI=0.1) for indicated time. Right: U2OS cells were infected with IAV^ΔNS1^ (MOI=1) for indicated time. **F.** Level of VSV N mRNA during infection. U2OS cells were infected with VSV^M51R^ (MOI=0.1 or MOI=1.0) for indicated time. **G.** Level of EMCV mRNA during infection. A549 cells were infected with EMCV (MOI=0.01 or 0.1) for indicated time. Data are presented in means ± SD. All data are representative of at least three independent experiments. *p* values were calculated using two-tailed unpaired Student’s t test (ns, *p*>0.05). Unless indicated otherwise, *p* values were calculated in comparison to WT values at equivalent conditions.

**Table S1.**
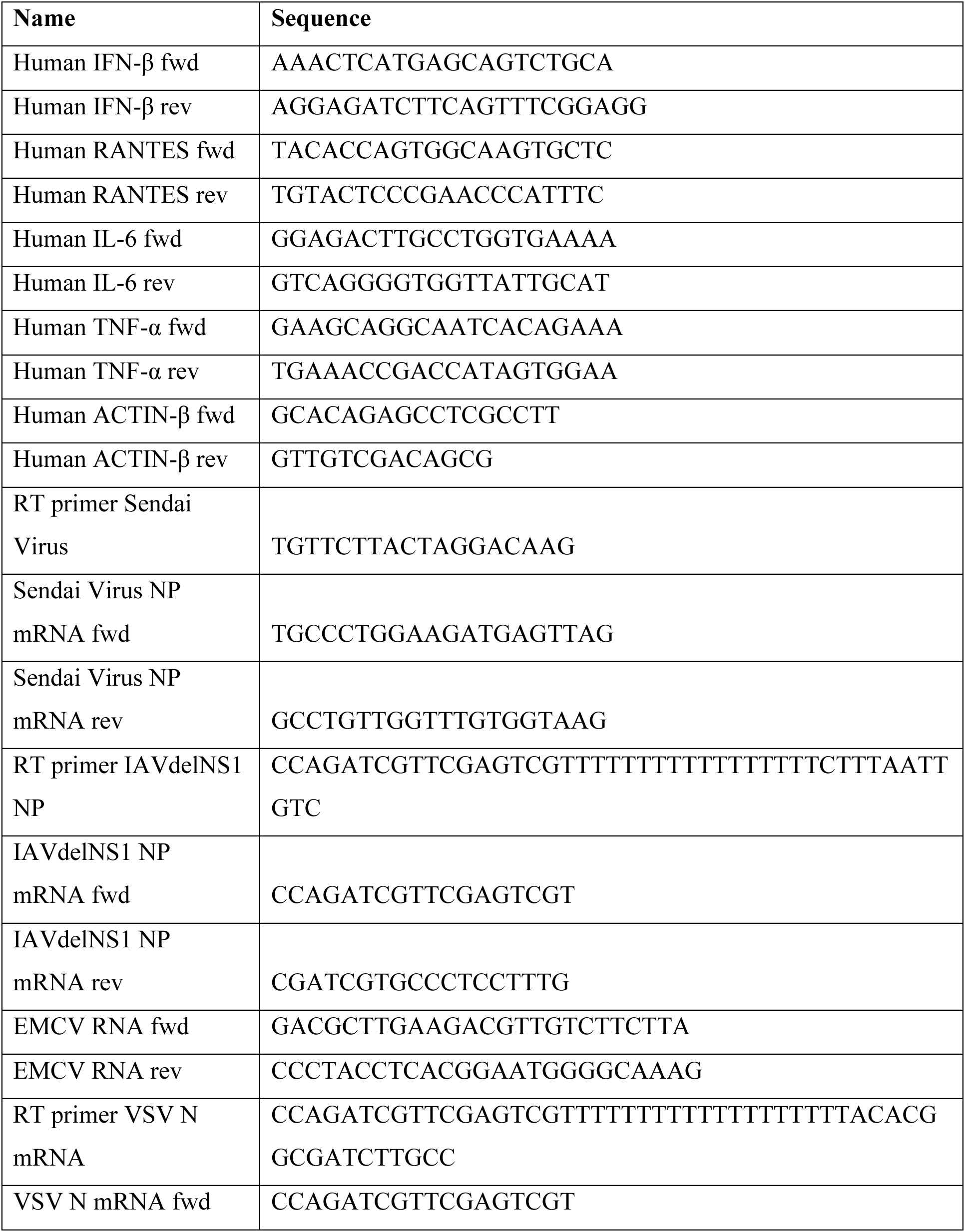
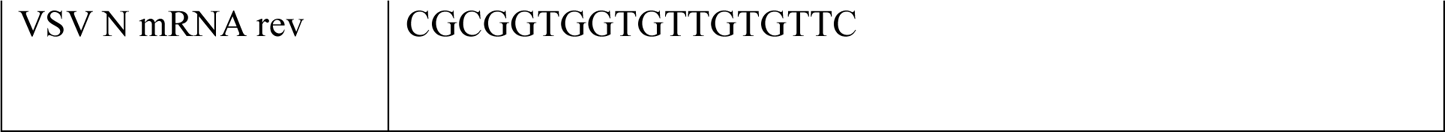
Primer Sequences.

**Table S2.**
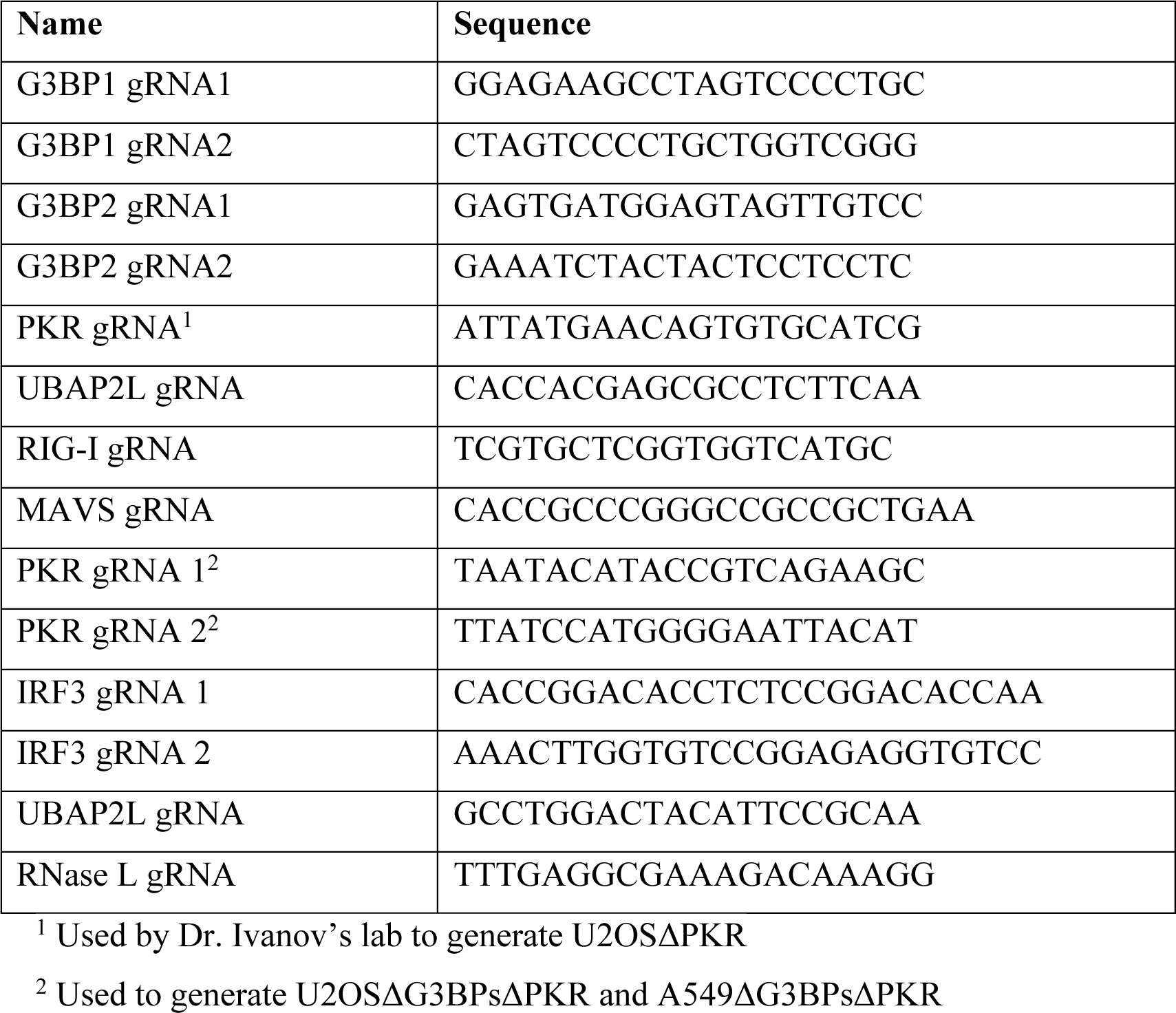
gRNA Sequences.

## Methods

### Material Preparation

#### Cell lines

U2OS, A549, HeLa and HEK293T cells were used for all experiments in this paper. The parental wild-type U2OS, G3BP1 and G3BP2 double knock-out cell lines were kindly provided by Dr. Paul J. Anderson and described elsewhere (Kedersha et al., 2016). The U2OS PKR knock-out cell line was generated by Dr. Shawn B. Lyons and kindly provided by him and by Dr. Paul J. Anderson (Aulas et al., 2017). The U2OS UBAP2L knock-out cell line was kindly provided by Dr. Paul J. Anderson (Sanders et al., 2020). Briefly, DNA encoding a guide RNA which targets the 5th exon of PKR were cloned into pCas-Guide (Origene) according to manufacturer’s instructions. U2OS cells were co-transfected with pCas-Guide-PKR and pDonor-D09 using Lipofectamine 2000. The following day, cells were selected with 1.5 mg/ml of puromycin for 48 hrs to select for transfectants. Single cell clones were isolated by limiting dilution and confirmed by western blotting and genomic sequencing. HEK293T cells were purchased from ATCC. The parental A549 cells were kindly provided by Dr. Susan Weiss, University of Pennsylvania (Li et al., 2017). The parental Hela cells were kindly provided by Dr. Gracjan Michlewski, University of Edinburgh.

For U2OS cells, for the generation of the PKR (in ΔG3BPs) knock out cell line, the ribonucleoprotein complex with gRNA and Alt-R® S.p. Cas9 Nuclease V3 was delivered using Lipofectamine 3000. After 48 hrs, media was refreshed, and single clones were isolated by limiting dilution and confirmed by western blotting and genomic sequencing. For the generation of the MAVS (both in WT and ΔG3BPs backgrounds), RNase L (in ΔG3BPs), RIG-I and IRF3 (in ΔG3BPs) knockout cell lines, gRNA was cloned in the pLentiCRISPRv2 vector using the restriction enzyme BsmBI. 293T cells were transfected with the following plasmids at a 3:1:0.7 ratio: (i) pLentiCRISPRv2 with the gRNA, (ii) pMD2.G VSV-G and (iii) psPAX2. U2OS (WT and ΔG3BPs) were infected with 0.45 µM filtered supernatants harvested at 48 hrs post-transfection for 48 hrs, then selected with 1 µg/ml neomycin.

For HelaΔG3BPs and A549ΔG3BPs, cells were first transduced using a gRNA targeting G3BP1 as described above. After transduction, the ribonucleoprotein complex with gRNA targeting G3BP2 and Alt-R® S.p. Cas9 Nuclease V3 (IDT) was delivered using Lipofectamine 3000, as described above as well. For generation of RNase L, MAVS and PKR knockout in A549 ΔG3BPs the same gRNA and same approach was used as described for the U2OS cells. A list of all gRNAs can be found in Supplementary Table 2. Single cell clones were isolated by limiting dilution and confirmed by western blotting and genomic sequencing. All cells were maintained at 5.0% CO2 in Dulbecco’s modified Eagle medium containing 10% fetal bovine serum and 1% L-glutamine. Cell lines were routinely tested for mycoplasma contamination.

#### Plasmids

The plentiCRISPRv2 puro was a gift from Brett Stringer (Addgene plasmid #98290). The plasmids psPAX2 and pMD2.G VSV-G were a kind gift from dr. James DeCaprio, MD, Dana-Farber Cancer institute. To generate the pInducer20-hG3BP1-IRES-G3BP2, hG3BP1 was cloned into pInducer20 (kindly provided by Dr. Hidde Ploegh, Boston Children’s Hospital) using NotI and AscI restriction sites followed by insertion of a SalI digested IRES-G3BP2 fragment generated by overlap PCR.

#### Viruses

Sendai virus (Cantell strain) was purchased from Charles River. Encephalomyocarditis virus (murine) was purchased from ATCC (VR-129B). VSV M51R and IAVdelNS1 were kindly provided by dr. Tristan Jordan and dr. Benjamin TenOever (NYU Langone Health). IAVdelNS1 was grown as previously described(Blanco-Melo et al., 2020). Briefly, MDCK cells expressing IAV-NS1 (MDCK-NS1 cells) were infected with IAVdelNS1 in EMEM containing 0.35% bovine serum albumin (BSA, MP Biomedicals), 4 mM L-glutamine, 10 mM HEPES, 0.15% NaHCO3, 1 μg/mL TPCK-trypsin (Sigma-Aldrich). Infectious titers were determined by TCID50/mL on MDCK-NS1 cells. VSV M51R was grown in in Vero cells in DMEM supplemented with 2% FBS. Viral supernatant was titered by plaque assay on naïve Vero cells. For both IAVdelNS1 and VSV-M51R stocks, viral supernatants were spun down to remove cellular debris. For SeV, EMCV, IAV^ΔNS1^ and VSV^M51R^ infection, A549 or U2OS cells were counted on the day of infection and subsequently infected with the listed MOI with 1 hr of absorption.

#### dsRNA

dsRNAs used in this study were prepared by in vitro T7 transcription as described previously (Peisley et al., 2012). The templates for RNA synthesis were generated by PCR amplification. The sequences of the 162 dsRNAs were taken from the first 150 bp of the MDA5 gene flanked by 5’-gggaga and 5’-tctccc. The two complementary strands were co-transcribed, and the duplex was separated from unannealed ssRNAs by 8.5% acrylamide gel electrophoresis in TBE buffer. RNA was gel-extracted using an Elutrap electroelution kit, ethanol precipitated, and stored in 20 mM Hepes, pH 7.0. Qualities of RNAs were analyzed by TBE polyacrylamide gel electrophoresis. For 3’-Cy5 labeling of RNA, the 3’end of RNA was oxidized with 0.1 M sodium meta-periodate (Pierce) overnight in 0.1 M NaOAc pH 5.4. The reaction was quenched with 250 mM KCl, buffer exchanged using Zeba desalting columns into 0.1 M NaOAc pH 5.4 and further incubated with Cy5-hydrazide for 6 hrs at RT.

#### RNA-seq

Cells were seeded in 6-wells plate and stimulated with 1 µg 162 bp dsRNA with 5’ ppp as described above. At indicated timepoints total RNAs were extracted from indicated cells using TRIzol reagent and RNA Clean & Concentrator. Quality control and mRNA-seq library construction were performed by Novogene Co. Libraries were sequenced on the Illumina NovaSeq 6000 instrument with a paired-end read length of 2 x 150 bp, which resulted in ∼20 M reads per sample. The raw sequence files were pre-processed using Trimmomatic v. 0.36 to trim Illumina adaptor sequences and low-quality bases. Trimmed reads were mapped to the human genome (UCSC hg38) using STAR aligner v. 2.5.4a. HTseq-count (v. 0.9.1) was used to count gene reads. Gene-count normalization and differential analysis were performed with DESeq25. Heatmaps were generated using Pheatmap. Scatter plots were generated using ggplot2.

#### RT-qPCR

Cells were transfected at 80% confluency with 500 ng/ml 162 bp dsRNA with 5’ppp. Lipofectamine 2000 was used for transfection with 2 µl lipofectamine reagent per µg of dsRNA diluted in 50 ul Opti-MEM per 500 ng of dsRNA. For mock transfection, cells were transfected with only lipofectamine reagent diluted in Opti-MEM. At indicated timepoints, total RNAs were extracted using TRIzol reagent and cDNA was synthesized using High-Capacity cDNA reverse transcription kit according to the manufacturer’s instruction. Real-time PCR was performed using a set of gene-specific primers or random primers, a SYBR Green Master Mix, and the StepOne™ Real-Time PCR Systems (Applied Biosystems). The full list of gene-specific primers can be found in table S1. For electroporation of dsRNA, 162 bp dsRNA with 5’ppp was electroporated into cells using a NucleofectorTM 2b (Lonza). 2 μg/ml of dsRNA was electroporated into cells using the cell line NucleofectorTM Kit L by following the manufacturer’s instructions using the U2OS program. At 8 hrs post-electroporation RNA was harvested for RT-qPCR. To determine the effect of TG on signaling, cells were treated with TG (1 μM) at 1 hr post-dsRNA transfection. In the G3BP1/2 complementation experiment, cells were seeded and expression of G3BP1-IRES-G3BP2 was induced with doxycycline for 24 hours and subsequently stimulated with dsRNA. To determine the effect of Q-VD-Oph on signaling, cells were pre-treated for 1 hr with either DMSO or 10 µM Q-VD-Oph prior to dsRNA transfection. For cGAMP stimulation, cells were permeabilized with digitonin buffer (50 mM HEPES pH 7.0, 100 mM KCl, 3 mM MgCl2, 0.1 mM DTT, 85 mM sucrose, 0.2% BSA, 1 mM ATP, 10 µg/ml digitonin with 10 or 20 μg/ml cGAMP for 30 min. After 30 min, complete Dulbecco’s modified Eagle medium with 10% fetal bovine serum was added and total RNA was extracted at 6 hrs post-transfection. For viral RNA measurement, virus-specific primers were used as listed in table S1.

#### ELISA

Cells were seeded in 6-well plates at 70% confluency. Cells were transfected with 162 bp dsRNA containing 5’ppp (500 ng/ml). At 6 hours post-transfection supernatant was harvested and used for ELISA. For IFN-β ELISA, the LumiKine™ Xpress hIFN-β 2.0 kit was used according to the manufacturer’s instructions. For RANTES, IL-6, TNF-α ELISA, the Human CCL5/RANTES Quantikine ELISA Kit, Human IL-6 Quantikine ELISA Kit, Human TNF-alpha Quantikine ELISA Kit were used respectively according to the manufacturer’s instructions. To determine the cytokine release upon SeV infection, cells were infected with 100 HA/mL SeV Cantell strain for 6 hours. Fresh supernatant was used for the ELISAs. The results were obtained using a Biotek M1 Synergy microplate reader using the manufacturer’s instructions.

#### Immunoblotting

Cells were seeded at 80% confluency in 6- or 12-well plates and transfected with 500 ng/ml 162 bp dsRNA with 5’ppp with lipofectamine 2000 as described above. At indicated timepoints, cells were lysed with 1% SDS lysis buffer (10 mM Tris pH 7.5, 150 mM NaCl, 10 mM DTT, 1% SDS), then boiled for 10 min. For the SUnSET assay, at 6 hrs post-transfection prior to puromycin (1 μg/mL) pulsing for 15 min. Proteins were resolved on 4–15% gradient gels, transferred to PVDF membranes, and blotted using standard procedures. Membranes were visualized using Amersham ECL reagent or SuperSignalTM West Femto Maximum Sensitivity Substrate.

#### Immunofluorescence microscopy and image analysis

U2OS, Hela or A549 cells were seeded on coverslips to reach 60-80% confluency the next day. Cells were stimulated with 500 ng/ml 162 bp dsRNA for 6 hrs as mentioned previously unless otherwise stated. At indicated timepoints, cells were fixed with 4% paraformaldehyde at RT for 10 min, and permeabilized with 0.2% Triton-X at RT for 10 min. Followed by blocking for 30 min at RT with 1% BSA in PBST, and staining using primary antibody for 1 hr at RT followed by 2 washes and then secondary antibody incubation for 1 hr. Hoechst 33342 was used to stain the nuclei. Images were obtained using a Zeiss Axio Imager M1 at 40X magnification. For immunofluorescence with Cy5-dsRNA, cells were transfected with 500 ng/mL of 162 bp dsRNA with 3’ Cy5 and 5’ ppp. For electroporation of dsRNA, 1 μg/mL of 162 bp dsRNA with 3’ Cy5 and 5’ppp was electroporated into cells using a NucleofectorTM 2b following the manufacturer’s instructions using the U2OS program. At 8 hrs post-electroporation the cells were fixed as mentioned before. For TG induction, cells were treated with 1 μM TG for 1 hrs prior to fixation. For SeV SG induction, cells were infected with 100 HA/mL SeV for 20 hrs prior to fixation. For SeV, VSV M51R and IAVdelNS1 protein level determination, cells were seeded for 90% confluency on the day of infection. Prior to infection cells were counted and infected. Cells were fixed at timepoints before cell death occurred. For ADAR1 knock-down, cells were transfected with pooled siRNA for ADAR1 or non-targeting control siRNA (50 μM). At 24 hrs post-trasfection media was changed to media containing 10 ng/mL recombinant human Interferon. At 48 hrs post-transfection, cells were fixed. For G3BP1-G3BP2 complementation, expression was induced for 24 hrs prior to 162 bp dsRNA transfection. For Q-VD-Oph treatment, cells were pre-treated with DMSO or Q-VD-Oph (10 µM) for 1 hr prior to dsRNA stimulation. All images were processed and analyzed with ImageJ. To highlight SGs, contrast adjustment was based on the mock transfected cells. To determine the Pearson Correlation Coefficient, the ImageJ plugin JaCOP was used on 10 separate FOV with at least 40 granules per FOV.

For nuclear IRF3 localization, U2OS WT and ΔG3BPs cells were stimulated with 162 bp dsRNA containing a 5’ppp (100 ng) for 6 or 16 hrs. Cells were prepared for immunofluorescence and stained with IRF3 (provided by Takashi Fujita) or DAPI. Images were taken randomly across the slide and the presence of IRF3 in the nucleus of each cell was quantified using ImageJ. The pixel intensity of nuclear IRF3 signal in each cell (a.u) were used to plot the data points.

For stress granule size analysis, U2OS WT, U2OSΔPKR and U2OSΔUBAP2L cells were stimulated with 162 bp dsRNA with 5’ppp for 6 hrs or 1 μM TG as mentioned previously. Cells were fixed and stained for G3BP1. Z-stack images (0.15 µM step size) were obtained using a Nikon TI2 motorized inverted microscope. All images were taken with a 60x oil-immersion lens. Stress granule size was determined by using the 3D Object Counter plugin in ImageJ. At least 200 SG from multiple fields of view (FOV) were picked randomly and analyzed. The percentage of cells that contained SG was determined by dividing the number of cells containing SG by the total amount of cells as measured by Hoechst 33342 staining for at least 5 FOV.

#### Cell death analysis and caspase cleavage assay

Cell detachment upon stimulation of cells was assessed at the indicated timepoints using the Nikon Eclipse TS2R at 20X magnification. For brightfield images, cells were seeded at 90% confluency in 12-well plates and transfected with 500 ng per well of 162 bp dsRNA with 5’ppp. At indicated timepoints, brightfield images were acquired from 3 different wells for each sample in duplicate. For the brightfield images of figure 4E, cells were stimulated with different cell death stimuli. U2OSΔG3BPs cells were transfected with 162 bp dsRNA (500 ng/ml) or treated with etoposide (20 µM) or Z-IETD-FMK (50 µM) plus TNFα (20 ng/ml) for 24 hrs. At 24 hrs post-treatment, the brightfield images were obtained using the Nikon Eclipse TS2R at 40X magnification.

For quantification of cell death, U2OS, Hela and A549 cells were seeded in 12-well plates and transfected with 500 ng per well with 162 bp dsRNA with 5’ppp unless otherwise indicated. At the indicated timepoints, cells were incubated with Sytox Green Nucleic Acid stain (2 µM final) and Hoechst 33342 (3,000-fold dilution) for 30 min. Sytox Green signal was measured with the Nikon Eclipse TS2R at 20X magnification using a 470 ex filter set. The percentage of dead cells was calculated as the number of Sytox positive cells divided by the total number of Hoechst positive cells using ImageJ. To determine the effect of different reagents on dsRNA-induced cell death, cells were pre-treated with DMSO, disulfiram (10 or 25 µM), Q-VD-Oph (10 µM), Human TNF-α Neutralizing Rabbit mAb (10, 100, 1000 ng/ml), Ruxolitinib (0.5 or 5 µM) or BX-795 (0.5 or 5 µM) for 1 hr prior to dsRNA transfection. For SeV, VSV M51R, IAVdelNS1 or EMCV infection, cells were counted on the day of infection and infected with the indicated MOI.

Live/Dead TM cell imaging kit was used to quantify cell death in U2OS WT and U2OSΔG3BP1/2 at the indicated timepoints. The manufacturer’s instructions were followed for this analysis. To quantify caspase 3/7 cleavage, cells were stimulated with dsRNA as previously mentioned and caspase 3/7 activity was analyzed using the CellEventTM Caspase 3-7 Green detection reagent. Similar to Sytox, the cells were stained with the Green detection reagent (1:30,000). The caspase 3/7 activity in figure 4B was measured using a Biotek M1 Synergy microplate reader using the manufacturer’s instructions. All experiments were performed at n=3.

#### IRF3 dimerization assay

This assay was adapted from the method described previously (Ahmad et al., 2018). Briefly, U2OS cells (mock or 112 bp dsRNA-transfected) were homogenized in hypotonic buffer (10 mM Tris pH 7.5, 10 mM KCl, 0.5 mM EGTA, 1.5 mM MgCl2, 1 mM sodium orthovanadate, 1X mammalian Protease Arrest) and centrifuged at 1,000 g for 5 min to pellet the nuclei. The supernatant (S1), containing the cytosolic and the mitochondrial fractions, was further centrifuged at 5,000 g for 15 min to pellet the crude mitochondrial fraction (P5). The P5 fraction was further washed once with isotonic buffer (hypotonic + 0.25 M D-Mannitol). The cytosolic fraction for the experiment was extracted from wild type untransfected U2OS cells using the same procedure as above except that the final spin was done at 18,000 g for 15 min and the supernatant S18 containing the cytosolic fraction was recovered. Subsequently, 35 μg of each P5 pellet was resuspended in 25 μl S18 (3 mg/ml) and used for IRF3 dimerization assay and Western blot analysis. 35S-IRF3 was prepared by in vitro translation using TnT T7 Coupled Reticulocyte Lysate System according to manufacturer’s instructions. The IRF3 dimerization was carried out by adding 16 μl (P5+S18) mix to 2 μl 35S-IRF3 in (20 mM HEPES pH 7.4, 4 mM MgCl2 and 2 mM ATP) in a total reaction volume of 20 μl. The reaction was incubated at 30°C for 1 h followed by centrifugation at 18,000 g for 5 min and the supernatant was subjected to native PAGE analysis. IRF3 dimerization was visualized by autoradiography and phosphorimaging on Amersham Typhoon 5 Biomolecular Imager. The image was quantified using ImageQuant.

#### RNase L activity analysis

Total RNA was isolated from cells and loaded on an RNA pico chip using an Agilent Bioanalyzer.

#### ADAR1 siRNA knock-down

Cells were seeded at 60% confluency and transfected with either pooled ADAR1 siRNA or non-targeting control siRNA (50 μM). After 24 hours, media was changed and recombinant human IFN-β was at 1 ng/mL or 10 ng/mL for 24 hours. After 48 hours, RNA was harvested and RT-qPCR and WB were performed to confirm ADAR1 knock-down.

#### Quantification and Statistical analysis

Average values and standard deviations were calculated using Microsoft excel and SPSS (IBM). The values for n represent biological replicates for cellular experiments or individual samples for biochemical assays. For each figure, individual replicate values were plotted together with the average values. The number of replicates is also indicated in the figure legends. Unless otherwise mentioned, all assays were performed in at least 3 independent experiments. p values were calculated using the two-tailed unpaired Student’s t test and are shown in the graphs.

